# Neurotoxicity of Propylene Glycol Butyl Ether: Multiomic Evidence from Human BrainSpheres

**DOI:** 10.1101/2025.05.22.655411

**Authors:** David Lopez-Rodriguez, David Pamies, Julien Boccard, Isabel Meister, Mathieu Galmiche, Tatjana Sajic, Estelle Maret, Noéline Héritier, Massimo Frangiamone, Aurélien Thomas, Nancy B. Hopf, Serge Rudaz, Marie-Gabrielle Zurich

## Abstract

Exposure to solvents may contribute to the development of neurodevelopmental and neurodegenerative diseases. Glycol ethers consist in a widely used class of organic solvents leading to workers and consumers exposure via many different applications. Ethylene glycol ethers are gradually being replaced by propylene glycol ethers thought to be less toxic. However, their neurotoxicity is not systematically assessed prior to placing them on the market. Therefore, this study investigated the potential neurotoxicity of propylene glycol butyl ether (PGBE) for which no official occupational limit has been established. To this aim, new approach methodologies have been used. Human induced pluripotent stem cells-derived BrainSpheres model was exposed to PGBE and to its main metabolite, 2-butoxypropanoic acid (2BPA). An integrative multiomic approach (transcriptomics, proteomics, metabolomics and lipidomics) was adopted to assess molecular alterations, derive benchmark concentrations and define potential mechanisms of action. PGBE was neurotoxic at occupationally relevant exposure concentrations. This was shown for the first time in human cells. And, although PGBE was more cytotoxic than 2BPA, both compounds showed very similar neurotoxicity. PGBE and 2BPA strongly affected the cell cycle, induced oxidative stress and perturbed energy and lipid metabolism. They also targeted specific nervous system processes, such as axon guidance and synapse organization. Finally, 2BPA may trigger ferroptosis by increased iron uptake. Our results show an urgent need for public health authorities to carefully assess the risk glycol ethers pose to humans, to properly protect the workers as well as individuals in the general population unknowingly exposed from indoor air contaminations.

## 1. INTRODUCTION

Organic solvents comprise a diverse class of chemicals that are widely present in numerous commercial products, contributing to their omnipresence in everyday life. Exposure to these organic solvents can occur in both workplace and residential environments. Much of the initial work on organic solvent toxicity originated in Scandinavia, where a neurobehavioral syndrome in painters, later termed “Painter’s syndrome”, was first described as leading to early retirement (Arlien-Soborg et al., 1979; Gregersen et al., 1978). Painter’s syndrome is characterized by symptoms such as memory impairment, mood disturbances, and reduced cognitive performance. While the solvents initially attributed to this syndrome, such as turpentine, thinner, acetone, toluene, benzene (Bates et al., 2016; Berr et al., 2010; Firestone and Gospe, 2009; Meyer-Baron et al., 2008) have since been largely replaced by other formulations, cognitive deficits are still associated with occupational solvent exposure. However, it remains unclear which solvents may be responsible for these effects today, as chemicals are not systematically assessed for neurotoxicity before being commercially available. Furthermore, the EU Classification, Labelling and Packaging (EU CLP) Regulation does not even include a classification for neurotoxicity.

Solvents target multiple brain regions. Chronic exposure may lead to global cognitive impairment, including deficits in memory, attention, energy, and personality, which are well described forms of dementia (Filley et al., 1990; Golbabaei et al., 2018; Sabbath et al., 2014). Once absorbed, solvents are preferentially distributed in lipid-rich structures, notably the adipose tissue and the brain, where they are mainly found in the white matter (Firestone and Gospe, 2009). It was originally hypothesized that solvents exert their toxic effects largely through nonspecific modulations of the membrane fluidity, and perturb the hydrophobic force regulating macromolecular interactions (Goldstein, 1984). They have also been shown to induce lipid peroxidation, leading to mitochondrial dysfunction, failure of electron transport and energy production (Fukumori et al., 2013; Revilla et al., 2007). However, evidence also supports that solvents interact with neurotransmitter receptors (Balster, 1998; Bowen et al., 2006).

Glycol ethers consist in a widely used class of solvents, leading to potential worker and consumer exposure via many different applications. They are found in the composition of paints, varnishes, engine fuels, hydraulic fuels, inks, cleaning products, pharmaceutical and cosmetic care, among others. The two main groups of glycol ethers (GEs) are based on ethylene glycol ethers (EGEs) and called the E-series, and on propylene glycol ethers (PGEs), called the P-series. Some members of the E-series have been shown to be reprotoxic, hematotoxic and neurotoxic in rats (Pomierny et al., 2013). Their toxic effects are largely thought to be due to their main metabolites, the alkoxyacetic acids, derived from their primary alcohol group (Aasmoe et al., 1998; Ghanayem et al., 1987; Miller et al., 1984). PGEs have two isomers, containing either a secondary (α-isomer) or a primary (β-isomer) alcohol. PGE α-isomers (>95% of the product) are metabolized in propylene glycol or conjugated with sulfate or glucuronide that are non-toxic (Reale et al., 2023). For this reason, PGEs are progressively replacing EGEs. However, the metabolism of the primary alcohol of PGE β-isomer is similar to EGEs and produces alkoxypropionic acids that may trigger toxicity. Exposure to propylene glycol methyl ether (PGME) has been shown to cause sedation in zebrafish, rats and rabbits (Miller et al., 1984; Puligilla et al., 2025; Spencer et al., 2002), similar to effects reported for other well-known organic solvents (Dick, 2006; Evans and Balster, 1991). PGME was also shown to cause headaches in humans exposed to vapour (Stewart et al., 1970). In a recent article using 3D rat brain primary cell cultures, propylene glycol butyl ether (PGBE) was found to be more neurotoxic than both PGME and EGME (Reale et al., 2023), the latter was banned due to repro- and hematotoxicity. Although based only on a few studies, the evidence consistently suggests that PGEs are associated with neurotoxicity, which remains largely unexplored.

The development of new approach methodologies (NAMs) and their use for risk assessment follows the 3R principles of replacing, reducing, and refining standard animal experiments (Russel and Burch, 1959). NAMs include high-throughput screening (HTS) bioassays, the field of omics applications, cell cultures, machine learning models and artificial intelligence (AI), quantitative structure–activity relationship (QSAR) predictions, and read-across (Marx et al., 2020; Rovida et al., 2020; Schmeisser et al., 2023; Sewell et al., 2024). Human induced pluripotent stem cells (hiPSCs)-derived 3D BrainSpheres is a NAM comprised of neurons, astrocytes and oligodendrocytes, and exhibiting spontaneous electrical activity (Carstens et al., 2025; Pamies et al., 2017). This model is well suited for neurotoxicological investigations (Nunes et al., 2023; Nunes et al., 2022; Pamies et al., 2018; Zhong et al., 2020). From a toxicological perspective, “omics” technologies, including genomics, proteomics, and metabolomics, are powerful approaches for generating relevant information on cellular changes induced by chemicals linked to adverse outcomes, and to detect early cellular responses underlying target organ toxicity. Recent workshops of the European Centre for Ecotoxicology and Toxicology of Chemicals (ECETOC) (Buesen et al., 2017) concluded that omics readouts could significantly contribute to risk assessment including the classification of substances, the mode of action of chemicals, and the identification of species-specific effects (Sauer et al., 2017). Due to the key roles of lipids in cellular processes (i.e, compartmentalization, signaling, and energy homeostasis), lipidomic analyses are now rapidly growing among -omics sciences and gained value for toxicology studies (Guardia-Escote et al., 2023; Tong et al., 2022). While single omics allow the investigation of a subset of the components of a particular biological process, multiomic approaches result in a more comprehensive picture, from gene to metabolites. Therefore, multi-omics data integration can considerably improve the confidence in identifying response pathways (Canzler et al., 2020; Shi et al., 2024).

In this study, we hypothesize that propylene glycol butyl ether (PGBE) is neurotoxic, and that its mechanisms of action can be accurately characterized with a combination of NAMs. The human BrainSpheres model was exposed to PGBE or its main metabolite 2-butoxypropanoic acid (2BPA). Then, an integrative multiomic approach (transcriptomics, proteomics, metabolomics and lipidomics) was used to assess molecular alterations, derive benchmark concentrations (BMC) at which responses are elicited and define potential mechanisms. Our results show the neurotoxicity of PGBE and urge public health authorities to carefully assess the risk glycol ethers pose to humans in order to properly protect the population.

## 2. MATERIAL AND METHODS

### 2.1 Experimental design

A multiomic approach was used to study the neurotoxicity of solvents in BrainSpheres (BSs), a 3D human iPSC-derived *in vitro* model. To capture the different levels of information from cellular and molecular pathways that may be impacted by organic solvents, transcriptomic, proteomic, metabolomic and lipidomic profiles were evaluated in BSs after a single (24/48h) or a repeated (1 week) exposure to PGBE or its metabolite 2BPA (Figure 1A). Transcriptomic analysis was performed after 24 h of exposure, while the remaining omics were evaluated after 48 h to account for differences in the timing of transcription, translation, and downstream processes. For each compound, four concentrations were evaluated: 0, 5, 10 and 20 mM. The workflow comprised bioinformatic approaches to characterize the multiomic profile of BSs and evaluate the neurotoxicity of solvents (Figure 1B).

**Figure 1.**
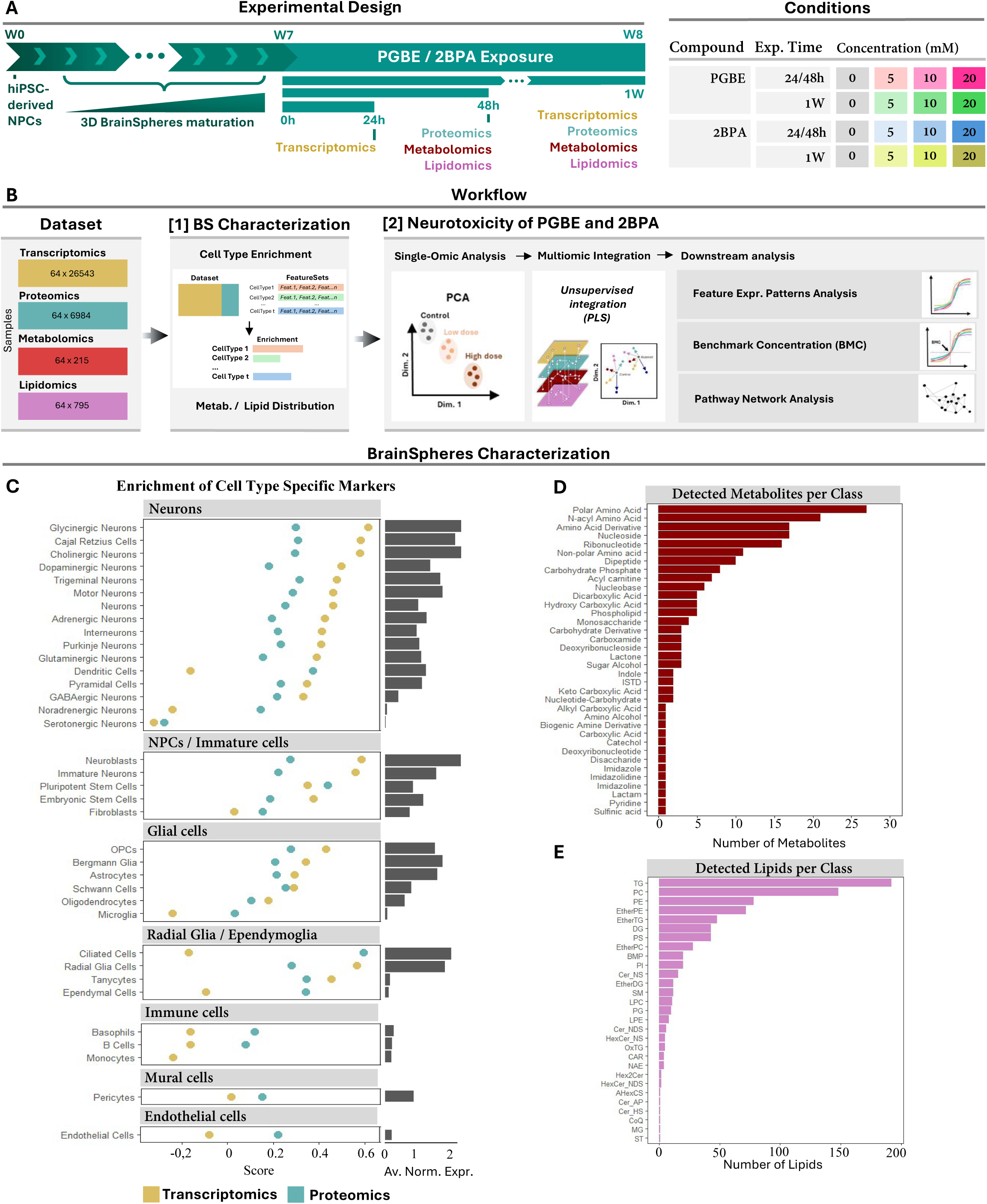
Experimental design, multiomic workflow and BS characterization. (**A**) hiPSC-derived 3D BrainSpheres (BS) were matured for seven weeks and exposed to PGBE or its metabolite 2BPA for either short-term (24h, 48h) or long-term repeated (1 week) durations at four different concentrations (0, 5, 10, or 20 mM). Multiomic experiments (i.e. transcriptomics, proteomics, metabolomics, lipidomics) were performed with four replicates per condition. (**B**) Multiomics workflow. First, BS characterization was conducted using cell-type enrichment and metabolite/lipid distribution analyses in control samples. Second, the neurotoxicity of PGBE and 2BPA was assessed through single-omic analyses (PCA and differential expression analysis), multiomic integration (PLS), and downstream multiomic analysis, including features expression pattern evaluation, benchmark concentration estimation, and pathway network analysis. (**C**) Dot plot showing the enrichment of cell-type-specific markers in control samples based on transcriptomics and proteomics data. Enrichment is represented by single sample gene set enrichment analysis (ssGSEA) scores, while bar graphs display the average normalized expression of gene sets associated with each evaluated cell type. (**D**) Bar plot illustrating the number of detected lipids and metabolites grouped by class.

### 2.2 BrainSpheres preparation

Neural progenitor cells (NPCs) were generated through embryoid body formation from induced pluripotent stem cells (iPSCs), themselves derived from human CCD-1079Sk fibroblasts (ATCC® CRL-2097) using Epstein–Barr virus-based vectors, as previously described (Wen et al., 2014). NPCs were then grown and amplified in cell culture flasks coated with Geltrex® (Thermo Fisher Scientific) in Neural Expansion Medium (NEM: 45% Neurobasal® Medium, 45% Advanced™ DMEM/F-12 Medium and 10% Neural Induction Supplement, Thermo Fisher Scientific). NPCs were kept at 37°C with 5% CO_2_. Medium was replaced every two or three days. Cell passages were performed at 90% confluency. BSs were then prepared from the NPCs as previously described (Pamies et al., 2017). At 95 % of NPCs confluency, cells were washed with Dulbecco’s Phosphate Buffered Saline (DPBS) and detached from the flasks using StemPro Accutase Cell Dissociation Reagent (GIBCO) for 2-3 min. 2×10^6^ cells per well were plated in 2 mL of NEM in non-treated six-well plates. The plates were placed in an incubator at 37°C and 5% CO_2_ under constant gyratory shaking (Kuhner orbital plate shaker, 88 rpm) allowing cell aggregation. After two days, cell medium was changed to differentiation medium consisting of Neurobasal Electro Medium, 1% Glutamax (Gibco), 2% B27/Neurobasal Electro, 0.01 μg/mL BDNF (PeproTech), 0.01 μg/mL GDNF (PeproTech) and 1% Penicillin-Streptomycin (10,000 U/mL). Medium changes with NDM were effectuated three times a week and cultures were kept until 8 weeks.

### 2.3 Preparation of chemicals

BSs were exposed to propylene glycol butyl ether (PGBE, CAS 5131-66-8, Merck, purity 99%) and to its metabolite 2-butoxypropionic acid (2BPA, CAS 14620-87-2, Merck, purity 95%). Solutions from the manufacturer were diluted in the differentiation cell culture medium to reach the final concentrations. The pH of the 2BPA solution from the manufacturer was very acidic and therefore was adjusted to 7.0 ± 0.5 before dilution in the medium. Seven-week-old BSs were exposed either one time and collected after 24 h or 48 h (24/48h) (acute exposure) or exposed repeatedly at each medium change, i.e., three times, during one week (1W) and collected at the end of the week, to 5, 10 or 20 mM of PGBE and 2BPA (Figure 1A). Untreated BSs were used as controls.

### 2.4 Cytotoxicity assay

Cytotoxicity of PGBE and 2BPA was measured after 1W repeated exposure to a range of concentrations of the chemicals (0 to 150 mM). Resazurin diluted in PBS was added directly to each well to reach a final concentration of 0.1 mg/mL. BSs were incubated for 3 h at 37°C, 5% CO_2_. Medium (100 µL) was then transferred in a 96-well plate. The fluorescent product was measured at 530 nm/590 ex/em with a Cytation 3 plate reader (BioTek). After subtraction of the background (resazurin solution only), results were expressed as percentage of untreated control cultures.

### 2.5 Lipidomics and Metabolomics

Lipidomics and metabolomics analysis were achieved with untargeted LC-HRMS approaches, as detailed in the Supplementary Material & Methods. Briefly, polar metabolites were first extracted from BSs by ultrasound-assisted lysis and extraction in a methanol / water (8:2, v/v) mixture. Then, lipids were extracted from the pellet of the first extraction in 100 % isopropanol. Lipid separation by UHPLC was achieved in reversed-phase mode with an Acquity Premier BEH C18 column (2.1 x 100 mm, 1.7 µm, Waters Corp.). Polar metabolites were separated on a zwitterionic Hilic column (Premier BEH zHilic, 2.1 x 100 mm, 1.7 µm, Waters Corp.) at pH 9.2. MS acquisition was performed in data-dependent mode on an Exploris 120 Mass Spectrometer (Thermo Fisher Scientific Inc.), equipped with a heated electrospray (HESI) source, in positive polarity for lipidomics and both positive and negative polarities for metabolomics. Data pre-processing and feature annotation was performed in MS-DIAL software (v.4.70). MS drift was corrected using LOESS algorithm and sample-to-sample variation in biological material by Probabilistic Quotient Normalization (PQN).

### 2.6 Proteomics

For proteomic analysis, approximately 40 mg of cell pellets from BSs were lysed in urea buffer, and protein denaturation, reduction, and alkylation were performed prior to overnight proteome digestion to peptides. The proteome digests were purified using C18 columns and spiked with Indexed retention time (iRT) peptides (RT-kit WR, Biognosys) before being analyzed by LC-MS. The samples were recorded on a Q-Exactive HF Hybrid Quadrupole-Orbitrap Mass Spectrometer (Thermo Fisher Scientific) operating in Data-Independent Acquisition MS (DIA-MS) mode. Data-dependent acquisition (DDA-MS) mode of respective sample digests was used for generation of sample-specific spectral library. The raw data files were processed with Spectronaut software (version:14.8.201029.47784, Biognosys) using workflow with spectral library, and quantitative protein matrices were generated as previously described (Wiskott et al., 2023). We identified more than 6.000 protein at 1% of false discovery rate (FDR) controlled at protein level data. Further details on the methodology, including sample preparation, MS parameters, and data processing, are provided in the Supplementary Material & Methods.

### 2.7 Transcriptomics

Extraction of total RNA was carried out using the Quiagen RNeasy Mini Kit. RNA concentration and integrity was measured using a fragment analyzer. Libraries were prepared following the BRB-seq protocol previously described (Alpern et al., 2019) and sequenced on a AVITI machine (Element Bioscience). Pre-processing was done in python (v3.9.6) following the NEXTFLOW pipeline for BRB-seq. Transcriptomic data was then analyzed in R (v4.4.0) using DESeq2 (v.1.44.0). Raw counts were imported and converted into a DeSeq2 object using the *DESeqDataSetFromMatrix*() function. Dispersion and size factor estimates were calculated, and data was fitted into a Negative Binomial GLM distribution using the *DESeq*() function.

### 2.8 Single omic bioinformatic analysis

To study data variance, a separate bioinformatic analysis of each omic dataset was performed. Dimensionality reduction was calculated following the normalization steps described for each omic dataset (Supplementary Material & Methods). Principal component analysis (PCA) was first done for all datasets using the prcomp() function from Stats (v.4.4.0). Differential analysis was then performed based on multiple t-test comparison methods based on *limma* with Benjamini-Hochberg (BH) correction. For transcriptomics, the Wald test was used to identify pairwise differentially expressed genes between treated and their respective control conditions using the Deseq2 function results().

### 2.9 Cell type enrichment analysis

A subset of transcriptomics and proteomics dataset containing exclusively non-exposed BSs was analysed to determine the enrichment of cell type specific markers present in 7-8 weeks-old BSs. A gene set enrichment analysis (GSEA) was applied to infer the relative cell type composition of BSs based on the enrichment of cell-type specific markers in the transcriptomics and proteomics dataset. GSEA generates a score per cell type considering the expression of markers in each sample and dataset. To allow the comparison between datasets, the data were first normalized and scaled. Then, to quantify the enrichment the single sample gene set enrichment analysis (ssGSEA) described in (Barbie et al., 2009) using the GSVA R package (v1.46) was used. ssGSEAis a non-parametric method that calculates a gene set enrichment score per sample as the normalized difference in empirical cumulative distribution functions (CDFs) of gene expression ranks inside and outside the gene set. As an input list, the cell type specific marker database of PanglaoDB, obtained from EnrichR libraries (*PanglaoDB_Augmented_2021*) was used.

### 2.10 Multiomic data integration and analysis

Following data normalization of each data source, as described in each corresponding section, the data was scaled according to their respective conditions. Features that displayed more than 50% of missing values were removed. Due to experimental limitations, different replicates were collected from different experiments for each omic dataset. As horizontal multiblock data integration requires multiple datasets measured in the same samples (shared sample mode), replicates were averaged to obtain a single value per experimental condition. For each block, rows represent condition means and columns are built from the features of each omic dataset. Finally, a supervised dimensionality reduction technique to assess shared and specific trends of variability between the data blocks was applied for multiomic data sources integration. This method allowed us to investigate potential similarities and/or differences between observations from the different conditions across the various omic datasets. Specifically, a horizontal multiblock integration using a Partial Least Square (PLS) model was applied using the R *mixOmics* package (v6.28) (Rohart et al., 2017). PLS maximizes the covariance between datasets (blocks) and generates latent variables that capture their shared variations. The *block*.*pls*() function was used using regression as a model and near-zero variance filtering and the transcriptomics dataset was used as *y*. Following integration, features displaying the same concentration-dependent patterns were searched. Hierarchical clustering was performed using the *<u>dist</u>*() and *hclust*() functions, using “*euclidean*” and “*ward.D2*” methods respectively. Clusters containing features with consistently increasing/decreasing expression pattern were selected for downstream analysis. The selected lists of features from each omic dataset were separated to perform functional enrichment analysis (for transcriptomics and proteomics) or to define the frequency by classes (for metabolomics and lipidomics). Functional enrichment analysis was performed with enrichR (version 3.1) using the most recent databases for “*Biological Process (2023)”.* Multiomic integration, clustering and downstream analysis was performed in R.

### 2.11 Benchmark concentration (BMC) analysis

To calculate the BMC for the multiomic datasets, we used the DRomics package (v2.6) in R, following these steps: (1) multiomic data matrices were normalized and scaled (prior to replicate averaging) for each condition (2BPA 24/48h, 2BPA 1W, PGBE 24/48h, PGBE 1W). (2) significantly responding features were selected using the ANOVA function and an FDR of 0.05 with the *itemselect*() function. This step allowed the reduction of noisy concentration-response signals before further processing. (3) Selected features were fitted to obtain the best-fitting concentration-response model using the *drcfit*() function. This function fits each concentration-response to a wide variety of monotonic and biphasic curves and selects the best fitting model based on the AIC (second-order Akaike criterion). Here, the focus was made on increasing/decreasing response curves as four concentrations were not sufficient to accurately define U-shape and bell-shape curves. (4) Calculation of the BMC was done according to the BMC-zSD methods proposed by EFSA (Committee et al., 2022). The BMC-zSD is the concentration at which the response is reaching the benchmark response (BMR), which is defined as: BMR=y0±z×SD. y0 represents the mean of the control response, z score is 1 by default (Larras et al., 2018), and SD is the residual standard deviation of the concentration-response fitted model calculated in the previous step. (5) Following BMC estimation, empirical cumulative density functions (ECDF) were calculated per experimental condition, where features were organized cumulatively according to their obtained BMC. These values were then used to evaluate the average cumulative BMC (acBMC) per condition, the median BMC per condition. (6) Functional enrichment analysis was performed in features from the transcriptomics and proteomics dataset having BMC below the acBMC as described in the previous section. BMC derived from the experimental cytotoxicity data was calculated following steps 3 and 4.

### 2.12 Multiomic pathway analysis

PaintOmics4 web tool (https://paintomics.uv.es/) was used to evaluate the pathways impacted by PGBE and 2BPA exposure using our multiomic dataset. This tool allows mapping multiple omics data to Kegg and Reactome pathways. PaintOmics4 requires a matrix of fold change comparisons per feature and a list of selected features of interest. To do so, the log fold change (logFC) of each treated concentration against its respective control for each of the four evaluated conditions (2BPA 24/48h, 2BPA 1W, PGBE 24/48h, PGBE 1W) was calculated. A reduced feature list was done, including only features differentially expressed and those having a BMC below the acBMC, calculated as described in previous sections. The software calculates enriched signaling pathways based on the fraction of features of interest overlapping pathway features and performing a Fisher exact test. To be restrictive and obtain the most affected pathways we selected only pathways that contained at least 75% of components affected by the exposure.

## 3. RESULTS

### 3.1 Multiomic Characterization of BrainSpheres

Similar cell type enrichment patterns found in the transcriptomic and proteomic datasets allowed the characterization of the cell types, metabolite and lipid composition of BSs. BSs strongly expressed markers associated with mature neuronal populations (i.e., Neurofilament Light Chain, (*NEFL*), Neurofilament Medium Chain (*NEFM*) and Tubulin Beta 3 Class III (*TUBB3*)), including glycinergic, cholinergic (Acetylcholinesterase (*ACHE*) and Choline Acetyltransferase (*CHAT*)), glutamatergic (Solute Carrier Family 17 Member 6 (*SLC17A6*), also called Vesicular Glutamate Transporter (*VGLUT2*)), dopaminergic (Tyrosine Hydroxylase (*TH*)) and GABAergic neurons (Glutamate Decarboxylase 1 and 2 (*GAD1* and *GAD2*)) (Figure 1C, Table S1). However, similar protein and gene expression levels of markers associated with immature neurons, such as doublecortin (*DCX*) and nestin (*NES*) were also found. Glial cells are also present in BSs, as indicated by the expression of cell-type specific markers for mature astrocytes, such as Solute Carrier Family 1 Member 2 and 3 (*SLC1A2 and SLC1A3*), both glutamate transporters (also called Glutamate Transporter-1 (*GLT1*) and Glutamate/Aspartate Transporter 1 (*GLAST1*), respectively) and to a lesser extent for mature oligodendrocytes (Proteolipid Protein 1 (*PLP1*) and 2’,3’-Cyclic Nucleotide 3’ Phosphodiesterase (*CNP*)) and oligodendrocyte precursor cells (SRY-Box Transcription Factor 10 (*SOX10*)). Markers for tanycytes, pericytes and endothelial cells were also observed. Finally, a few pluripotent cells were still undifferentiated in 8-week-old BSs, as indicated by the presence of SRY-Box Transcription Factor 2 (*SOX2*).

The metabolomic data block showed 215 metabolites in BSs (Figure 1D, Table S1). Polar amino acids (27, e.g., glutamic acid, tyrosine) and N-acyl amino acids (21) are highly represented classes, followed by amino acid derivatives (17, e.g., creatine, phosphocreatine, glutathione) and nucleosides (17), ribonucleotides (16, e.g., AMP, cAMP, ADP and ATP), non-polar amino acids (11, e.g., tryptophane), dipeptides (10) and carbohydrate phosphates (8, e.g., gructose-1,6-diphosphate, glucose-6-phosphate, glycerol-3-phosphate). The presence of neurotransmitters as gamma aminobutyric acid (GABA), dopamine and serotonin were also observed.

The lipidomic assay reported 795 annotated lipids in BSs (Figure 1E, Table S1), among those 192 triacylglycerol species (TG), 148 phosphatidylcholines (PC), 78 phosphatidylethanolamines (PE), 72 ether phosphatidylethanolamines (EtherPE), and 48 ether triacylglycerols (EtherTG). Sphingomyelins (SM), abundant lipids in cell membranes were also present (12).

Overall, the multiomic characterization suggests that 8-week-old BSs are highly enriched in markers associated to the main brain cell types, neurons, astrocytes and oligodendrocytes that are found in immature and advanced maturation stages. Furthermore, lipids and metabolites associated with brain cell populations and functions could be identified and in particular, neurotransmitters and their precursors.

### 3.2 Multiomic data integration and differential expression analysis

A multiomic data integration analysis was performed to evaluate the neurotoxicity of PGBE and 2BPA and gain insight into their mechanisms of action. As experiments for PGBE and 2BPA exposures were performed in separate batches and they exhibited distinct profiles (Figure S1A), multiomic integration was performed on each compound separately to explore the differences between exposure times and concentrations for each chemical.

Multiomic integration of the PGBE exposure dataset revealed a concentration-dependent effect after 1W of repeated exposure (separation along dimension 1) (Figure 2A). After 24/48h, PGBE-exposed samples were also clearly separated from their controls (along dimension 2), but no strong concentration-dependent effect could be observed. Dimensions 1 and 2 in Figure 2A explain 33% and 24% of the variance, respectively. All omics contributed relatively equally to the variance of dimension 1, whereas the lipidomic dataset contributed more importantly than the others to the second dimension (Figure S1B). Single-omic analysis showed that the concentration-dependent effects of PGBE were mainly observed in the transcriptomic and lipidomic datasets, after both exposure durations (Figure S1C). The concentration-dependent PGBE effects was less marked in the proteomics and metabolomics datasets.

**Figure 2.**
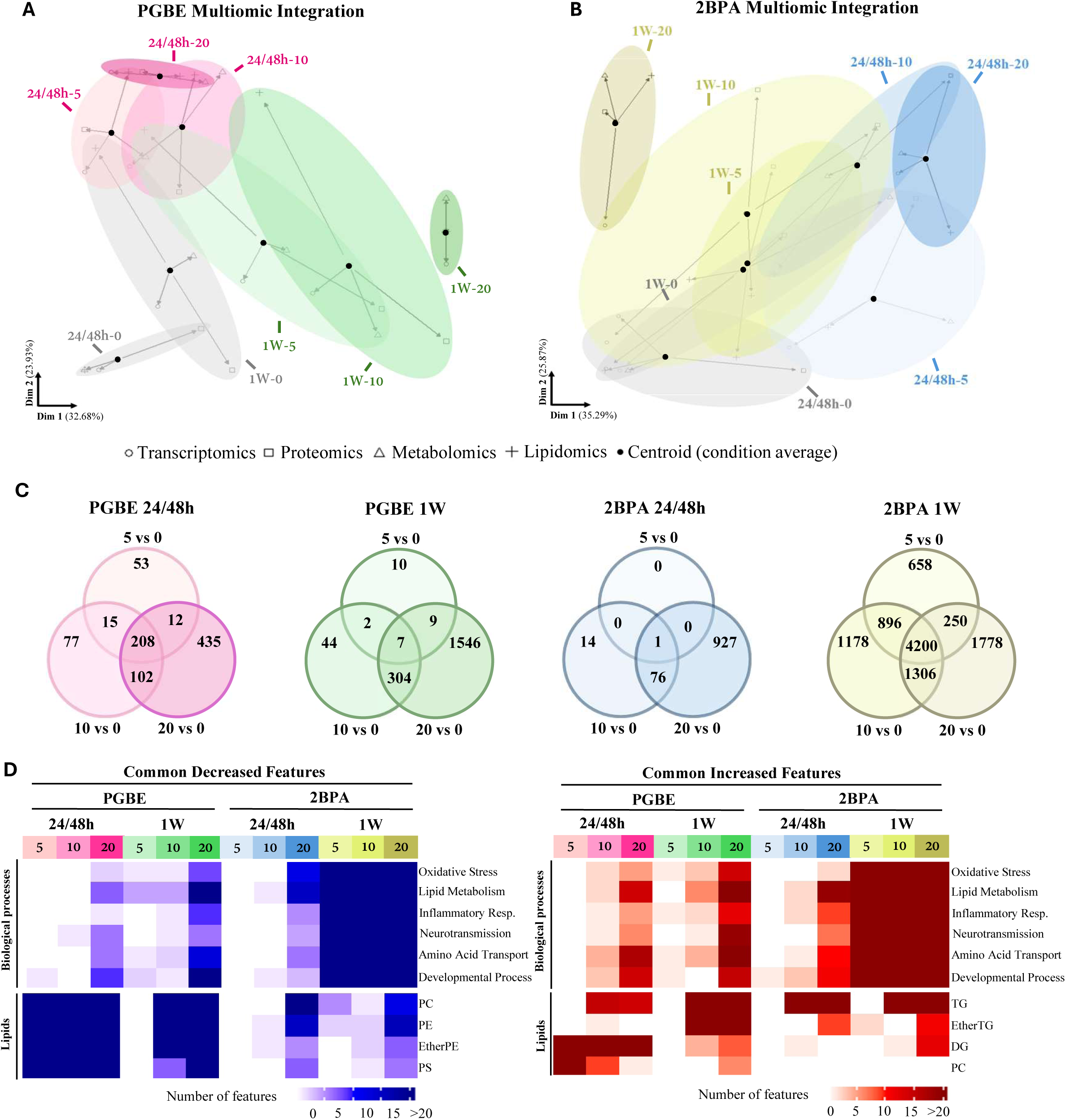
Multiomic PLS model and differential expression analysis. (**A-B**) Arrow plot displaying the first two components from the multiomic PLS model for PGBE and 2BPA. Arrow starting points (black dots) represent centroids (average of replicates) per condition across all omics. The arrow heads indicate the corresponding condition averages in each individual omic dataset. Arrow lengths reflect the level of agreement or variance divergence between the multiomic model and individual omic datasets. (**C**) Venn diagram showing the number of differentially expressed features per conditions and concentrations. Values represent features from all omic datasets. (**D**) Heatmap illustrating the biological processes associated with the most commonly upregulated and downregulated features in the transcriptomics and proteomics datasets, along with the number of lipids belonging to the most affected lipid classes.

Multiomic integration of the 2BPA dataset showed a less clear concentration-dependent impact of the exposure (Figure 2B). The strongest differences were observed after both exposure durations to the highest concentration (20 mM). The first two dimensions of the PLS model explained between 35% and 26% of the variance and again lipidomics had a greater contribution to the second dimension as compared to the other datasets (Figure S1B). This second dimension was the most closely linked to the separation of experimental conditions based on exposure concentration. Single-omic analysis showed that the transcriptomic, lipidomic and to a lesser extent proteomic datasets are impacted by 2BPA exposure, particularly after 1W (Figure S1D).

A differential expression analysis (DEA) was computed using each individual omic dataset to highlight features that were significantly impacted by the exposure (Table S2). Globally, the number of modulated features increased according to concentration (Figure 2C, Figure S2). The 1W-repeated exposure to 2BPA displayed the highest number of differentially-expressed features.

Heatmaps show genes and proteins commonly decreased or increased by PGBE and 2BPA (Figure 2D). These features were associated with the following biological processes: oxidative stress (e.g., *HSP90B1, HSPA4L, HSPA5, SESN2*), lipid metabolism, inflammatory response, neurotransmission, amino acid transport (e.g., *SLC3A2, SLC7A5*) and developmental processes. The number of decreased or increased features was highest after 1 week of exposure to 2BPA. Additionally, phosphatidylcholines (PC), phosphatidylethanolamines (PE), ether phosphatidylethanolamines (EtherPE) and phosphatidylserines (PS) where among the most decreased lipids by both chemicals, whereas triacylglycerols (TG), ether triacylglycerols (EtherTG) and diacylglycerols (DG) were within the most increased ones (Figure 2D, Table S2). Other features were almost only impacted by 2BPA, such as transferrin receptor highly significantly increased at gene (*TFRC*) and protein (P02786) levels after single and repeated exposure, and its ligand transferrin (P02787), significantly upregulated after 1 week (Figure S2, Table S2).

Overall, both compounds share numerous differentially modulated expressed features involved in biological processes frequently encountered in neurotoxicity, i.e., oxidative stress, lipid metabolism and inflammation.

### 3.3 Concentration-response patterns of PGBE and 2BPA

Following multiomic data integration, features showing concentration-dependent increase or decrease were highlighted and further investigated separately for each compound and time-point (Figure 3A). This analysis allowed the identification of the most impacted biological functions, while considering the relationships between omic datasets and between the concentrations (Figures 3-4, Figures S3-S4).

**Figure 3.**
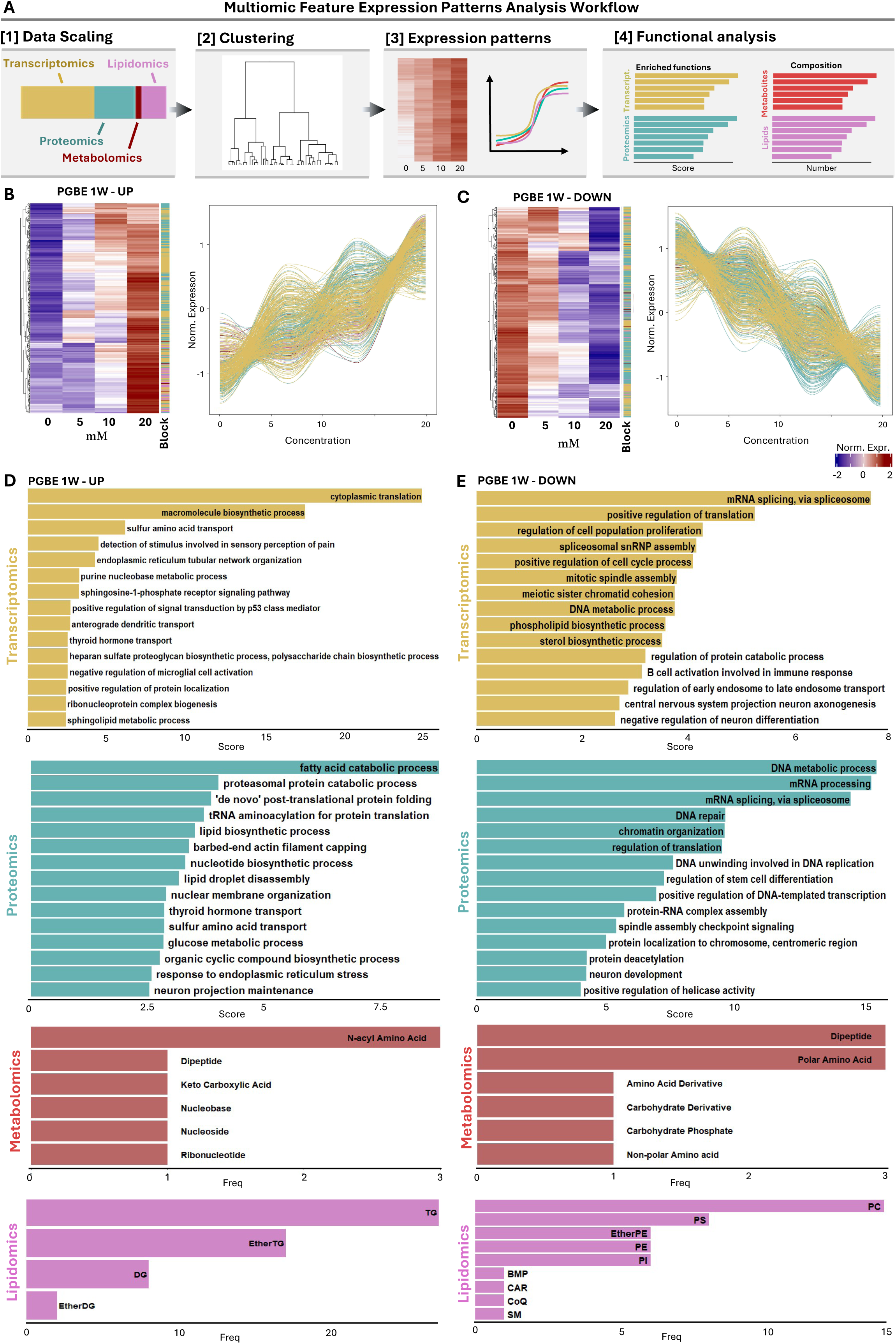
Multiomic feature pattern analysis. (**A**) Following dataset scaling, a hierarchical clustering was performed to identify specific concentration-response expression patterns. Downstream functional analysis was performed for transcriptomics and proteomics, while distribution of lipid and metabolite classes was evaluated for each identified pattern. (**B-C**) Heatmap and line graph showing features increasing or decreasing in a concentration-dependent manner in the PGBE 1W condition. (**D-E**) Enrichment analysis of biological processes associated with genes and proteins displaying increased or decreased expression patterns in the PGBE 1W condition. Scores represent the enrichment of gene ontology terms. The number of metabolites and lipids belonging to their respective classes is also shown.

**Figure 4.**
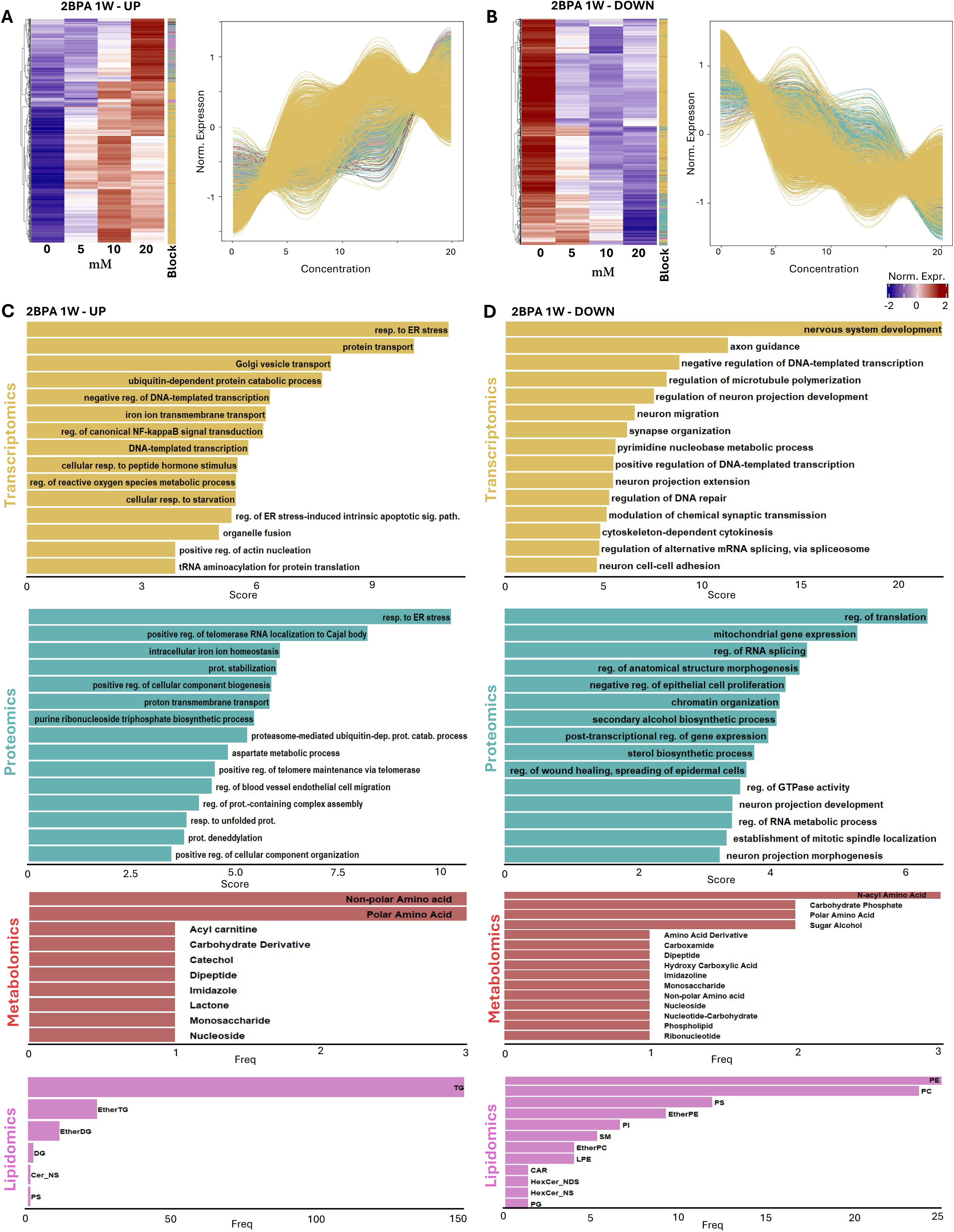
2BPA 1W Multiomic feature expression patterns. (**A-B**) Heatmap and line graph showing features that increase or decrease in a concentration-dependent manner in the 2BPA 1W condition. (**D-E**) Enrichment analysis of biological processes associated with genes and proteins displaying increased or decreased expression patterns in the 2BPA 1W condition. The distribution of metabolites and lipids is also shown.

Enrichment analysis revealed that PGBE significantly up- and down-regulated features involved in very general biological processes, in a concentration-dependent way (Figure 3 B-E, Figure S3 A-D). After both exposure durations these features were associated with cell cycle, cytoplasmic translation, synthesis of macromolecules, amino acid transport, as well as lipid and glucose metabolism. Furthermore, a few biological processes specific for the nervous system were deregulated after 1 week of exposure to PGBE, such as axonogenesis and projection maintenance. An accumulation of TGs, EtherTGs and DGs was observed after the single and the repeated exposure, whereas a decrease was observed for PCs, PEs and EtherPEs (Figure 3 D-E, Figure S3 C-D). Nucleic acid constituents were increased and carbohydrates decreased at both time-points (Figure 3 D-E, Figure S3 C-D).

The exposure to 2BPA also strongly impacted general biological processes such as mitosis, translation and energy metabolism (Figure 4 A-D and S4 A-D). However, the highest enrichment was found for the up-regulation of features associated with ER stress response, and the generation of ROS was also highlighted. Iron transport was also increased. Finally, 2BPA impacted a higher number of nervous system specific processes than PGBE. After the 1-week exposure, increased features were associated with nervous system development, axon guidance and synapse organization (Figure 4D).

Overall, these results suggest that PGBE and 2BPA share common actions on general biological processes, including cell cycle and energy metabolism. According to these concentration-response pattern analyses, the main differences in their action appear to be the higher number of nervous system-specific biological processes impacted by 2BPA and its ability to upregulate iron transport.

### 3.4 Benchmark concentrations (BMC)

To assess the concentrations at which PGBE and 2BPA trigger a neurotoxic response, benchmark concentrations (BMCs) were determined from the multiomic analysis (Figure 5A) and compared to the BMC determined from the cytotoxicity data. The cytotoxicity-derived BMC of PGBE and 2BPA was 25.6 mM (23.4-26.9) and 63.8 mM (52.8-72.3) respectively, indicating a higher cytotoxicity of PGBE compared to its metabolite 2BPA (Figure 5B).

**Figure 5.**
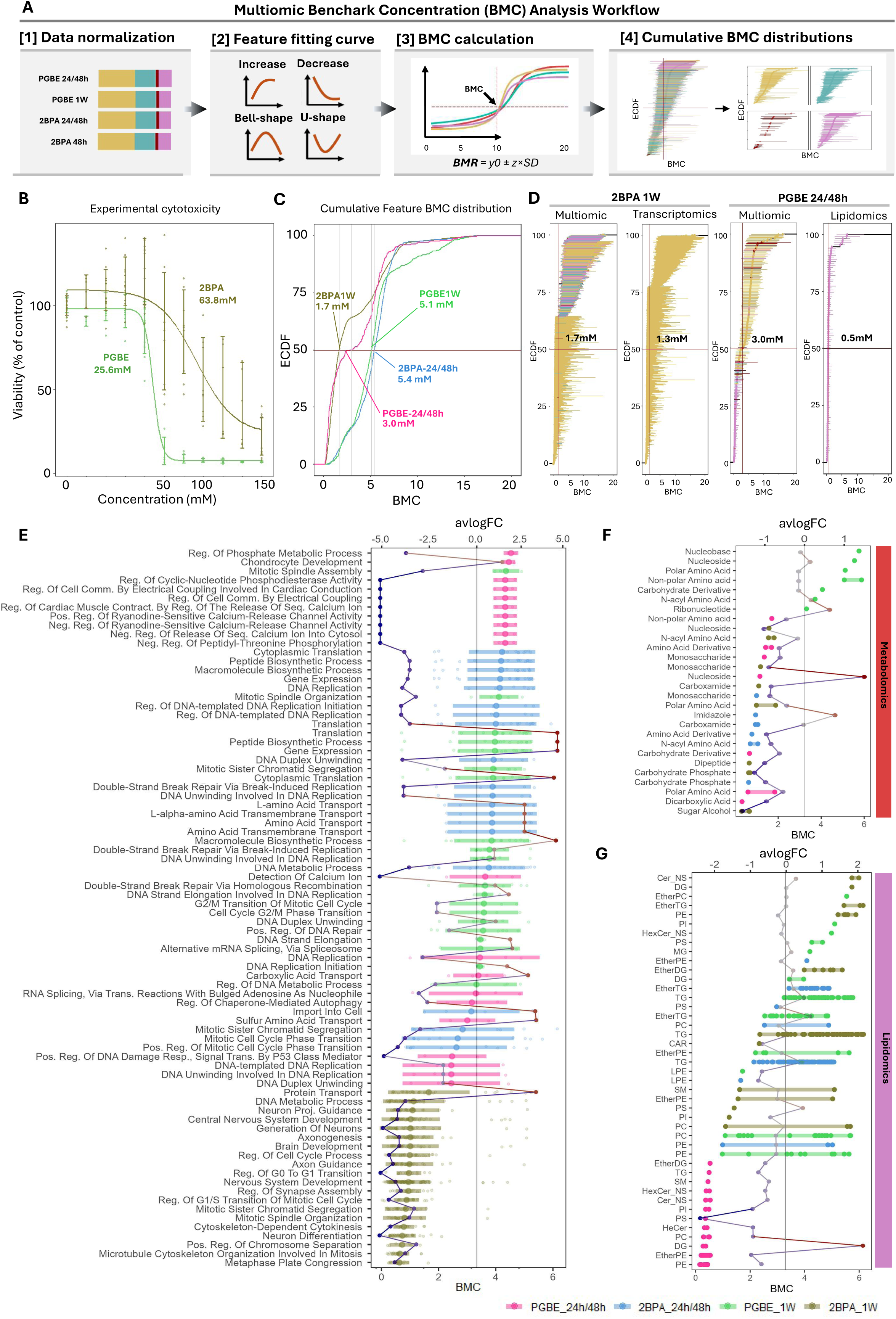
Multiomic BMC analysis. (**A**) Scaled and normalized multiomic data per condition was fitted to concentration-response curves. Fitted curves were used to calculate BMC cumulative distributions. (**B**) Fitted concentration-response curve for experimental cytotoxicity of PGBE and 2BPA using concentrations ranging from 0 to 150mM. (**C**) Multiomic BMC empirical cumulative distribution function (ECDF) per condition. Horizontal red line displays the median value of the distribution. (**D**) EDCF BMC for 2BPA 1W and PGBE 24/48h exposure, displaying BMC from multiomics, transcriptomics or lipidomic datasets. Vertical lines represent confident intervals. (**E**) Functional enrichment analysis of features below the avBMCs per condition. The BMC of genes and proteins within each pathway is displayed as a dotplot (bottom x axis), while average log fold change (avlogFC) of these features (5mM versus 0mM) is represented as a line plot (top x axis). For the BMC dotplot, central dots represent the average of gene and protein BMCs associated with the pathways. Horizontal lines represent confidence intervals. (**F**) Distribution of metabolite and lipid classes of features below the avBMC per condition. Average log fold change (avlogFC) of these features (5mM versus 0mM) is represented as a line plot (top x axis).

From the multiomic dataset, BMCs were derived for 24/48h and 1W of PGBE and 2BPA exposure. Given the large amount of data that can be obtained from multiomic datasets, the average cumulative BMC (acBMC) was considered as a reference value. The acBMC is the median of the computed significant BMC responses per condition. Results showed a lower acBMC for 2BPA compared to PGBE after 1 week of exposure (1.7 mM and 5.1 mM, respectively), whereas after 24/48h, it was the opposite (5.4 mM and 3.0 mM, for 2BPA and PGBE, respectively) (Figure 5C, Table S3). The acBMC for each omic separately showed that the low acBMC after 1-week 2BPA was driven by features from the transcriptomic dataset (1.3 mM), whereas lipidomics drove the acBMC for PGBE 24/48 h (0.5 mM) (Figure 5D). All other values are given in Figure S5A-D.

To discriminate the most sensitive biological functions affected by the solvents, a functional analysis was performed using only the features in the transcriptomic and proteomic datasets having a BMC lower than the acBMC value per omics for each condition. GO terms retrieved after exposure to PGBE (both durations) and 2BPA 24/48h were all associated with general biological functions, including cell cycle, translation, amino acid transport, peptide biosynthesis (Figure 5E). While 1 week of exposure to 2BPA was also associated with some cell cycle processes, it particularly decreased nervous system specific processes, including nervous system development, axon guidance, axonogenesis and synapse assembly (Figure 5E).

In the metabolomic dataset, features with the lowest BMC were found to be mostly decreased (Figure 5F). Specifically, the 1-week 2BPA exposure decreased sugar alcohol metabolites, carbohydrate phosphate, dipeptide, polar amino acids, monosaccharides, N-acyl amino acids and nucleosides, while 1W PGBE exposure impacted ribonucleotides, n-acyl amino acids cabohydrate derivatives, polar and non-polar amino acids, nucleosides and nucleobases, at higher concentrations. Finally, features with the lowest BMC in the lipidomic dataset (e.g., PEs, EtherPEs, DGs, PCs) (Figure 5G) were impacted by the 24/48h exposure to PGBE. All conditions appeared to impact PEs and TGs at different concentrations below the acBMC.

Altogether these results confirm the neurotoxicity of PGBE and its metabolite 2BPA (acBMC < cytotoxicity BMC), and that cell cycle is strongly affected by both compounds.

### 3.5 Multiomic pathway enrichment analysis

To gain deeper insights into the mechanisms of action of PGBE and 2BPA, multiomic pathway enrichment analysis was performed (Figure 6A). The pathways were then sorted to evidence the most significantly modified ones (Combined pValue(Fisher) ≤ 0.05), showing at least 75% of the total number of features modified and comprising at least 10 features. The number of pathways retrieved was higher after exposure to 2BPA (21 pathways for 1 week exposure, 18 after 48 h) than after PGBE (17 after 1 week, 4 after 48 h) (Table S4).

**Figure 6.**
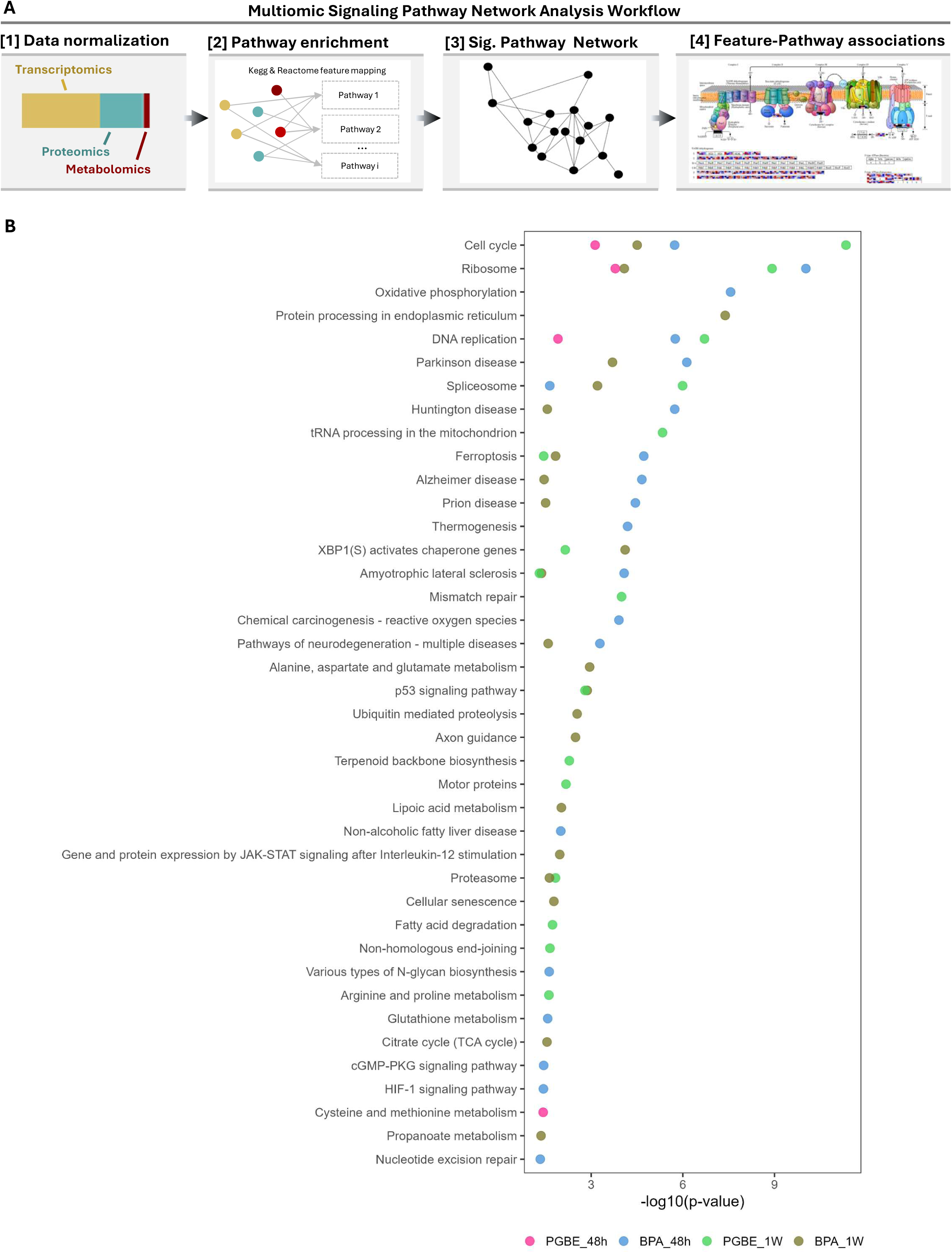
Multiomic pathway analysis. (**A**) Significant DEA features and features below the avBMC for the transcriptomic, proteomic and metabolomic datasets were mapped to Kegg and Reactome pathways using PaintOmics. (**B**) Plot displaying the Kegg and Reactome pathways for each condition with a p-value of > 0.05 and having at least 75% of their components impacted.

PGBE and 2BPA significantly deregulated cell cycle and ribosome pathways in the 4 conditions tested (Figure 6B, Table S4), suggesting they are early and important targets of these chemicals. After 1 week of exposure, PGBE and 2BPA also significantly impacted the same stress pathways (Figure 6B, Table S4), i.e., XBP1(S) activates chaperone genes, p53 signaling pathway and finally ferroptosis, an iron-dependent form of nonapoptotic cell death (Dixon et al., 2012). The latter was highlighted after both exposure durations to 2BPA, in line with the highly significant up-regulation of TRFC described under chapter 3.2. Both compounds were also associated with the amyotrophic lateral sclerosis disease (ALS) pathway.

Concerning pathways significantly affected only by one chemical, PGBE impacted fatty acid degradation, terpenoid backbone biosynthesis and amino acid (arginine and proline) metabolism after 1 week of exposure (Figure 6B, Table S4), whereas 2BPA was associated with perturbation of alanine, aspartate and glutamate metabolism, lipoic acid metabolism and several nervous system specific pathways, i.e., axon guidance and neurodegenerative disease pathways other than ALS. Further, 2BPA affected energy metabolism, as seen by deregulation of the citrate cycle after 1 week.

These results suggest that PGBE and 2BPA clearly act on the cell cycle, and that 2BPA targets more pathways specific to the nervous system.

## 4. DISCUSSION

PGBE is a glycol ether widely used in industrial products (De Luca and Hopf, 2025), however data on its neurotoxicity still remains scarce. In this study, NAMs based on human cells and high throughput omic technologies were used for their potential to improve the efficiency and reliability of chemical toxicity screening, as well as to decipher chemicals’ mechanisms of action. Omics approaches already proved to be very useful for various applications in the field of neurotoxicology (Olesti et al., 2021; Pamies et al., 2023; Schultz et al., 2015; Schvartz et al., 2019; Tobolkina et al., 2023). However, multiomic integration methods that may help to holistically identify altered pathways, are not commonly used. Here, the neurotoxicity and mechanisms of action of PGBE and its main metabolite 2BPA were investigated in a hiPSC-derived 3D brain *in vitro* model (BrainSpheres, BS) using a multiomic approach. The scenario modeled in this study was based on predicted brain concentration ranges in workers exposed to structurally similar propylene glycol ether. Consequently, the findings can be directly applied for risk assessment relevant for both occupational and public health. They can help in predicting the neurotoxicity of these widely used but yet poorly characterized family of chemicals (De Luca and Hopf, 2025).

Knowing that the development of a human brain does not finish before adulthood, the presence of immature cell type markers revealed by the multi-omics profiling of BSs was not surprising. And, very importantly, the presence of various subpopulations of mature neurons, such as glutamatergic, GABAergic, cholinergic and dopaminergic neurons, as well as astrocytes and oligodendrocytes was observed. The higher presence of neurons than glial cells was in line with previous results (Pamies et al., 2017). These organ-specific characteristics emphasized the interest of using BSs for the present neurotoxicological study.

Various BMCs were extrapolated in this study. One BMC for cytotoxicity was determined for chronic PGBE and 2BPA exposure, and BMCs from each feature of the multiomic dataset fitting concentration-dependent curves were computed, again for each chemical and time-point. PGBE was found to be more cytotoxic than its metabolite 2BPA. This is somewhat in contradiction with the common thinking that metabolites of glycol ethers are responsible for the toxicity of their parent compounds. This may be true for ethylene glycol ethers (Multigner et al., 2005), but not for propylene glycol ethers as shown here. The higher cytotoxicity of PGBE may be explained by its slightly higher blood-brain barrier permeability (Werner et al., 2025).

The average cumulative BMC derived from the multiomic datasets was lower than the cytotoxicity-derived BMC for PGBE and 2BPA (5 times and 38 times lower, respectively) after 1 week of repeated exposure, strongly suggesting the neurotoxic potential of these chemicals. The neurotoxic action of 2BPA was also emphasized by the enrichment in nervous system-specific pathways exhibiting BMCs below the acBMC value (5.4 mM), such as nervous system development, axon guidance and synapse assembly. These results are in line with the neurotoxic effects of PGBE already suggested after exposure of 3D rat primary mixed brain cell cultures (Reale et al., 2023), where effects on brain cell-type specific parameters were shown around 3.3 mM. This also suggests that rat and human species are similarly sensitive to PGBE.

BMC derivation is the preferred European Food Safety Authority (EFSA) approach for identifying a reference point that can be used as a starting point for risk assessment (Committee et al., 2022). BMC modeling of omic datasets is promising to characterize neurotoxicity as it defines features potentially impacted by very low concentrations. It also allows evaluating chemicals’ mechanisms of toxicity and supports the development of adverse outcome pathways (Gao et al., 2024; Li et al., 2022; Matteo et al., 2022; Vuong et al., 2025). Features displaying a low BMC indicate the existence of a sharp increased/decreased expression at the lowest experimental concentration evaluated. However, interpretation should be approached with caution due to the limited number of tested concentrations (Larras et al., 2018). The approach was strengthened by combining BMC analysis with gene ontology to uncover ensembles of features involved in specific functions, sharing expression patterns and similarly low BMCs. As powerful as this tool may be to identify the impact on cellular functions of very low concentrations, the results should be experimentally confirmed.

To identify pathways impacted by PGBE and 2BPA acute and repeated exposure, differentially-expressed features together with features displaying a BMC below the acBMC were selected. Strikingly, pathway analysis pointed to the cell cycle as being strongly inhibited by both chemicals in a concentration- and time-dependent manner. The p53 pathway was also strongly modified by both compounds after 1 week, suggesting DNA disruption and the initiation of cell cycle arrest, cellular senescence or apoptosis (Harris and Levine, 2005). This was confirmed by the downregulation of the mismatch repair pathway by PGBE.

The strong and early modifications in lipids observed after exposure to both chemicals may constitute a key event in the neurotoxicity of these compounds. Due to their physico-chemical properties, glycol ethers could disrupt lipid assemblies from the cell membrane and perturb its function. The plasma membrane contains microdomains enriched in certain glycosphingolipids, gangliosides, and cholesterol that form membrane/lipid rafts (MLRs), where many molecules, including signaling receptors and ion channels are anchored (Head et al., 2014; Schnaar et al., 2022). Several cytoskeletal components and enzymes that regulate the cytoskeleton localize to MLRs. PE-rich domains seem to stabilize cytoskeletal attachments and participate in endocytosis and exocytosis, which are tightly regulated by actin filament remodeling (Horn and Jaiswal, 2019; Kapus and Janmey, 2013; Schroer et al., 2020), while PC domains contribute to connect actin filaments to the plasma membrane (Czogalla and Sikorski, 2010; Ha and Exton, 1993; Innocenti, 2018; Schroer et al., 2020). Accordingly, by decreasing PEs and PCs, solvents may contribute to the downregulation of genes involved in cytoskeleton (e.g. RHOA, ACTB), microtubule organization (e.g. TACC3, MAPT) and intermediate filament (GFAP) observed in this study. This could in turn lead to altered cell shape, motility, and signal transduction, ultimately disturbing brain cell function and plasticity. The decrease of PEs, EtherPEs, PCs and SMs may also indicate a perturbation of the myelin sheath, since they constitute its principal lipid components (Kister and Kister, 2022; Montani, 2021; Poitelon et al., 2020), as already proposed for various other solvents (Rumsby and Finean, 1966).

The unfolded protein response (UPR) is mounted when endoplasmic reticulum (ER) stress is detected and serves primarily to return normal ER function (J. H. Lin and Lavail, 2010). A wide variety of chemicals are known to activate the UPR, including the N-linked glycosylation inhibitor tunicamycin, oxidizers such as dithiothreitol, sodium arsenite and cisplatin (Jennings et al., 2013). Ultimately, if the UPR fails to return the cell to homoeostasis, apoptosis is initiated. This pathway, in particular the XBP1 branch, was activated after the repeated exposure to both compounds, indicating that they induced ER stress.

Ferroptosis is a type of non-apoptotic regulated cell death involving intracellular iron toxicity and lipid peroxidation (Chen et al., 2021). Ferroptosis can be triggered by excessive iron accumulation or inhibition of glutathione peroxidase 4 (GPX4) (Chen et al., 2021), a key enzyme in the antioxidant defense. In this study, 2BPA highly significantly induced the upregulation of transferrin (TF) and its receptor (TFRC) at the mRNA and protein levels, suggesting higher iron uptake. This clearly could promote radical oxygen species (ROS) production and lipid peroxidation (Dixon et al., 2012). Ferritin (FTL) and ferritin receptor subunits (Ferritin Light Chain, (FTL), Ferritin Heavy Chain 1 (FTH1)) mRNA levels were also increased after 2BPA, likely as a counter mechanism to capture free iron by binding to transferrin, and then extrude it from the cells through the ferritin receptor. Altogether, these results suggest that 2BPA interferes with iron transport that may lead to increased lipid peroxidation and ultimately to cell death by ferroptosis. A high percentage of features of this pathway were also modified by PGBE, however, the main components of iron uptake, TFRC and TF were not regulated, indicating that PGBE unlikely triggers higher iron uptake.

Besides pathways commonly deregulated by both compounds, enrichment analysis also indicated that 2BPA exerted specific effects on nervous system related pathways, such as axon guidance. 2BPA decreased the expression of the guidance cues secreted in the extracellular matrix, as well as their specific receptors present on developing neurons. The binding of the guidance cues to their respective membrane receptors determines the direction of axon growth (Garbe and Bashaw, 2004; Huber et al., 2003). Dysfunction of axon guidance has been associated with the onset of neurodevelopmental and neurodegenerative disorders (L. Lin et al., 2009; McFadden and Minshew, 2013). Indeed, guidance cue systems continue to be expressed in the adult brain and are emerging as important mediators of synaptic plasticity and fine-tuning of mature neural networks (Mahmud et al., 2023). Furthermore, epidemiological studies suggest that prenatal exposure to ethylene glycol ethers is associated with altered brain activity and performance during a motor inhibition task in children (Binter et al., 2019) and with lower verbal comprehension score and visuospatial performance (Beranger et al., 2017). According to the results of the current study, the neurotoxicity of 2BPA may be at least partially attributed to its ability to perturb axon guidance.

Inhalation is the most important route of exposure for glycol ethers, and blood can absorb an extensive quantity of PGME and PGBE that is immediately bioavailable for highly vascularized tissues (Borgatta et al., 2022) where they may induce toxic effects. An in silico toxicokinetic (TK) model previously developed and calibrated (Reale et al., 2023), adapted to workers with high physical activity, predicts total brain PGME concentration above 4 mM, after 8 h of exposure to PGME 100 ppm at high physical activity, raising to above 6 mM after 5 consecutive working days. In the latter situation, free PGME in the brain is predicted to reach more than 2 mM. Since PGBE is much more lipophilic than PGME (LogP 1.15 and −0.49, respectively) and has a higher in vitro determined blood-brain barrier permeability (Werner et al., 2025), it is expected to reach even higher total and free brain concentrations in case of exposure to 100 ppm. The neurotoxic effects of PGBE observed in the present study occurred at concentrations comparable to those predicted to reach the human brain following exposure to PGME, a very similar propylene glycol ether. While no official occupational exposure limit (OEL) has been established for PGBE—likely due to the assumption of low toxicity and the absence of required neurotoxicity testing—this propylene glycol ether concentration (100 ppm) is known to cause symptoms such as headaches (Stewart et al., 1970). Despite PGBE being present in numerous commercial products for decades, workplace exposures are neither regulated nor routinely monitored. Our findings highlight the urgent need to establish OELs for PGBE, particularly in light of its widespread use and potential neurotoxic effects. PGBE solvents are also commonly found in consumer products used indoors such as cleaning agents, paints, and air fresheners, and thus, contribute to chronic low-level exposures. This is especially concerning for vulnerable populations, including children, the elderly, and individuals with pre-existing health conditions, who may be more susceptible to the neurotoxic effects of prolonged exposure, even at lower concentrations. Given the potential for cumulative health impacts in residential environments, our results highlight the need to evaluate neurotoxicity of chemicals within a broader public health context.

PGEs are sold as a mixture of 2 isomers, with the bulk having a secondary alcohol group (α-isomer) and primary alcohol group (β-isomer). The β-isomer is oxidized by alcohol dehydrogenase and aldehyde dehydrogenase to alkoxypropionic acid in the body (Aasmoe et al., 1998; Miller et al., 1984; Werner et al., 2025) as the major reaction product, thought to be responsible for the toxicity of glycol ethers (Aasmoe et al., 1998; Ghanayem et al., 1987; Miller et al., 1984). The neurotoxicity of PGBE described in this study may be partly due to 2BPA since BSs were recently shown to be able to metabolize propylene glycol ethers (Werner et al., 2025), however, our results show that the neurotoxicity of PGBE itself warrant further investigation and risk assessment by public health authorities.

## 5. CONCLUSION

In conclusion, our in vitro study provides, for the first time, strong evidence that PGBE is neurotoxic for human cells, at concentrations compatible with predicted concentrations to be reached in the brains of exposed workers. Furthermore, although 2BPA is less cytotoxic than PGBE, the neurotoxicity of both compounds is comparable. Our study also illustrates the benefit of using a multiomic approach to tackle mechanisms of neurotoxicity. Indeed, metabolomics and lipidomics, as phenotypic read-outs, are essential components in building and confirming pathways retrieved thanks to more commonly measured transcriptomics and proteomics. Finally, our study urges public health institutions to better characterize exposures and the consequent health risks associated with handling PGE containing products. PGEs can induce neurotoxic responses at concentrations relevant in occupational settings. Such effects may contribute to the onset of neurological disorders, particularly in workers. However, children may also be at risk, as a recent study (Garlantézec et al., 2020) showed the presence of glycol ethers in their urine likely due to the use of domestic products.

## CRediT authorship contribution statement

Conceptualization: MGZ, DLR and NBH; Data curation: IM, MG and TS; Formal analysis: DLR, IM, MG, TS and JB; Funding acquisition: NBH, SR and MGZ; Investigation: DP, IM, MG, TS and NH; Project administration: MGZ; Software: DLR and JB; Visualization: DLR; Writing - original draft: DLR, MF and MGZ; Writing - review & editing: All authors

## Acknowledgements

This work was in part supported by the Swiss Centre for Applied Human toxicology (SCAHT), Basel, Switzerland and the Federal Office of Public Health (FOPH), Bern, Switzerland.

