## Supplemental Materials for "Neurotoxicity of Propylene Glycol Butyl Ether: Multiomic Evidence from Human BrainSpheres"

**Table of Contents**


- **Supplementary methods**
- **Figure S1.** Single and multiomic models. (**A**) Arrow plot displaying the first two components from the multiomic PLS model for PGBE and 2BPA together. Arrow starting points (asterisks) represent centroids (average of replicates) per condition across all omics. The arrow endpoints indicate the corresponding condition averages in each individual omic dataset. Arrow lengths reflect the level of agreement or divergence between the multiomic model and individual omic datasets. (**B**) Bar plot displaying the explained variance contribution of each dataset of dimensions 1 to 7 for the PGBE and 2BPA models. (**C-D**) PCA displaying the first two components for each individual dataset.
- **Figure S2.** Differential expression analysis for the transcriptomics and proteomic datasets.
- **Figure S3.** PGBE 24/48h Multiomic feature expression patterns. (**A-B**) Heatmap and line graph showing features that increase or decrease in a concentration-dependent manner in the PGBE 24/48h condition. (**D-E**) Enrichment analysis of biological processes associated with genes and proteins displaying increased or decreased expression patterns in the PGBE 24/48h condition. The distribution of metabolites and lipids is also shown.
- **Figure S4.** 2BPA 24/48h Multiomic feature expression patterns. (**A-B**) Heatmap and line graph showing features that increase or decrease in a concentration-dependent manner in the 2BPA 24/48h condition. (**D-E**) Enrichment analysis of biological processes associated with genes and proteins displaying increased or decreased expression patterns in the 2BPA 24/48h condition. The distribution of metabolites and lipids is also shown.
- **Figure S5.** Multiomic BMC analysis. (**A-D**) Multiomic BMC empirical cumulative distribution function (ECDF) for PGBE 24-48h, PGBE 1W, 2BPA 24-48h and 2BPA 1W conditions. Vertical

red line displays the median value of the distribution and vertical lines represent confident intervals per feature.

- **Additional File. Excel document.**

**Supplementary methods**

**Lipidomics and metabolomics.**

1. Sample preparation

Brain Spheres in 1.5 mL Eppendorfs were first washed in 1 mL ammonium acetate (150 mM in water) to remove traces of culture medium. The wash solution was removed after gravity sedimentation of Brain Spheres and 1 mL methanol / water (8:2, v/v) containing 1 µM CHES (Sigma-Aldrich) as technical internal standard was added for polar metabolomics extraction. After a few seconds vortexing (Vortex Genie-2, Scientific Industries), spheres were disrupted in an ice bath by 10 consecutive cycles of ultrasonication of 10 seconds each separated by 30 second breaks to reduce heat production (Vibra-Cell VCX500, Sonics & Materials Inc.). Sonication intensity was set at 30 % amplitude. After extraction, samples were incubated at -20°C for 1 h to promote protein precipitation and centrifuged at 14000 g for 15 minutes at 4°C (5430R, Eppendorf SE). The supernatant containing polar metabolites was collected and evaporated to dryness on low temperature setting (Savant SC210A concentrator, Thermo Fisher Scientific Inc.). The pellet obtained after metabolite extraction was resuspended in 1 mL isopropanol containing 1 µM LPC 18:1-d7 (Sigma-Aldrich) as technical internal standard. Ultrasonication was performed similarly for 5 cycles at 45 % amplitude to re-homogenize the pellet. Samples were then shaken in a ThermoMixer (Eppendorf SE) for 15 minutes at 1200 rpm and 4°C and centrifuged at 14000 g for 15 minutes at 4°C. The supernatant was collected and evaporated to dryness.

On the day of analysis, dry extracts were reconstituted for metabolomics in 150 µL acetonitrile / water (7:3, v/v) containing 2 µM HEPES (Sigma-Aldrich) as technical internal standard, and for lipidomics in 250 µL methanol (100 %) containing 1 µM Cer 18:1-d7/15:0 (Sigma-Aldrich).

Recovery was enhanced by vortexing samples for 10 seconds and shaking them at 1200 rpm and 4°C for 15 minutes. Samples were then placed in the freezer overnight to trigger the precipitation of potential particles. After centrifugation (14000 g, 4°C, 15 min), 40 µL or 50µL of each sample was pooled for constituting QC samples for metabolomics or lipidomics, respectively. Polar metabolomics extracts were transferred into MaxQuant PP vials (Waters Corp.) and lipidomic extracts into TruView glass vials (Waters Corp.).

### 2. Metabolomics and lipidomics data acquistion

Analytical measurements were performed on a Vanquish Horizon UHPLC system (Thermo Fisher Scientific Inc.) hyphenated to an Exploris 120 Mass Spectrometer (Thermo Fisher Scientific Inc.) equipped with a heated electrospray (HESI) source.

Lipidomic analysis was performed with reversed-phased liquid chromatography (RPLC). The analytical column was an Acquity Premier BEH $C_{18}$ with dimensions 2.1 x 100 mm, 1.7 µm particle size and 130 Å pore size (Waters Corp.) equipped with a 5 mm guard column. Solvent A was acetonitrile / water (60:40, v/v) with 10 mM ammonium acetate. Solvent B was isopropanol / acetonitrile (90:10, v/v) with 10 mM ammonium acetate. All solvents were LC-MS grade (Fisher Optima). Mobile phase B was ramped up from 20% to 50% in 3 min, then up to 85% at 10 min, and to 97% 15 min. Mobile phase composition was then return to initial conditions for 4 minutes, resulting in a total run time of 19 min. The flow rate was 0.4 mL/min and the column temperature was set at 50°C. The injection volume was 2 µL. MS source parameters were the following: spray voltage = + 3.4 kV, transfer tube temperature = 340°C, sheath gas flow = 55 AU, auxiliary gas flow = 13 AU, sweep gas flow = 2 AU. $MS^1$ spectra were acquired as centroid data in full scan mode, from 220 to 1200 m/z, with 60k resolution at 200 m/z and AGC at 100%. $MS^2$ centroid data were acquired in data-dependent mode by fragmentation of the four most intense $MS^1$ singly charged ions (Top4). The intensity threshold was set at $2 \cdot 10^4$ for precursor fragmentation and there was a 2 second dynamic exclusion of previously fragmented precursor ions within a 10-ppm tolerance. The isolation window was set at 0.9 m/z, the collision energy at 20 eV, and the resolution was 15k at 200 m/z. An exclusion list was prepared using a solvent blank injection to exclude contaminants from the selection of $MS^1$ ions to be fragmented.

Polar metabolites were separated on a Premier BEH zHilic (2.1 x 100 mm, 1.7 µm, Waters Corp.) column protected with a 5 mm guard column. Solvent A was 20 mM ammonium acetate in water (100%) with pH adjusted to 9.2 with 0.1 % ammonium hydroxide solution (25%, Fisher Scientific). Solvent B was 100% acetonitrile. The elution of polar metabolites was obtained by increasing percentage of solvent A from 10 to 60 % from 0.3 to 12.3 min). The column was washed at 100 % A for 4.7 column volumes and reconditioned at 10 % A for 9.2 column volumes for a total run time of 21 min. The flow rate at elution was 0.4 mL/min, the column temperature was 30°C and the injection volume was 4 µL. MS acquisition was done in negative mode, with spray voltage set at -2.3 kV, the transfer tube temperature at 320°C, the sheath gas flow at 50 AU, the auxiliary gas flow at 10 AU, the sweep gas flow at 2 AU. Full scan $MS^1$ centroid spectra were acquired with 60k resolution at 200 m/z and AGC 100%, in a mass range from 80 to 600 m/z and the $MS^2$ data-dependent spectra with the Top4 approach. The intensity threshold was set at $1 \cdot 10^5$ for precursor fragmentation and there was a 3 second dynamic exclusion of previously fragmented precursor ions within a 10-ppm tolerance. The isolation window was set at 0.9 m/z, the collision energy at 15 eV, and the resolution was 15k at 200 m/z. An exclusion list was prepared using a solvent blank injection to remove contaminants from the $MS^1$ ions to be fragmented.

### 3. Data pre-processing and annotation

MS-DIAL (v.4.70) was used for feature annotation, both for lipidomics and metabolomics datasets (Tsugawa et al., 2015). Peak picking was performed with a minimum peak height of 200 and a mass slice width of 0.1. Peaks were aligned across all chromatograms with a 0.1 min retention time tolerance and a 0.01 m/z $MS^1$ tolerance.

Features measured with the lipidomic analysis protocol were annotated according to *in silico* MS/MS matches with the lipid database embedded in MS-DIAL. The specified criteria for peak picking were a maximal 0.01 Da difference in accurate mass (for both $MS^1$ and $MS^2$ levels) and a minimal $1 \cdot 10^3$ signal intensity. Annotated lipids exhibiting poor peak shapes, abnormally high mass difference (> 5 ppm deviation from theoretical mass) or low signal-to-noise (i.e. below 3) were excluded. Only one adduct per lipid species was retained, and the only adducts represented in the curated dataset were $[M+H]^+$, $[M+H-H_2O]^+$ and $[M+NH_4]^+$. The m/z vs. RT visualization of MS-DIAL was used to verify that the retention of annotated features followed the equivalent carbon rule commonly accepted for lipids (Ovčačíková et al., 2016).

Features from polar metabolomics were first matched against an in-house database comprising values of retention time (RT) and accurate mass (AM). A total of 122 compounds were annotated at level 1. Additional annotations in the dataset were performed at levels 2 and 3, on features exhibiting a high intensity and signal-to-noise ratio, with accurate mass ($MS^1$) and fragmentation pattern ($MS^2$) matches against HMDB-indexed compounds using MS-FINDER (v.3.52 ) (Tsugawa et al., 2016).

4. Data filtering and correcting

Both lipidomics and metabolomics datasets were filtered and corrected before integration in the multi-omic analysis pipeline. The instrumental drift was corrected with an R-based LOESS regression algorithm (Cleveland, 1979, Dunn et al., 2011) . After drift correction, features displaying a CV higher than 20% on pooled QCs and/or a D-ratio higher than 40 were excluded from the dataset. Additionally, features with more than 33% missingness (defined as peak area below the limit of detection, i.e. 3 times the peak area measured in the blank) in samples, and/or more than 10% of missingness in pooled QCs, were also excluded. Finally, to account for differences in total cellular material in the samples, Probabilistic Quotient Normalization (PQN) was performed on both metabolomics and lipidomics using an in-house platform.

**Proteomics.** Approximately 40 mg of cell pellets from BSs were lysed in the 300 µL of 8M UREA buffer. Lysis was performed using Tissue Lyser II (Qiagen) with the addition of 4 tungsten carbide beads 3mm, for a period of 3 minutes at 20s-1 frequency. Lysed samples were centrifuged at 2 500 rpm for 2 minutes and 200 µL of supernatant was collected in a new tube and stored at -80ºC. Prior to proteome digestion to peptides, samples were thawed on ice and denaturation was performed in 8M UREA buffer followed by standard process of protein reduction and alkylation each time for 1 hour at controlled temperature of 37ºC in thermomixer. We used 10mM tris(carboxyethyl)phosphine (Sigma-Aldrich) for protein reduction and 20mM iodoacetamide (Sigma-Aldrich) for alkylation. The urea concentration was then adjusted to 1 M by diluting the samples with 50 mM AMBIC buffer, giving a total of 1450µL of solution for protein digestion. Proteome digestion was performed overnight at 37°C using 25µL of sequencing grade porcine trypsin (Promega) per sample, and digestion was stopped with 10µL of formic acid (FA). To remove undigested proteome and sample contaminants, the collected peptides were purified on C18 silica MicroSpin columns (The Nest Group, Inc.) by low-speed centrifugation (i.e., 400g for 2 min) and five consecutive wash steps in 0.1% aqueous formic acid (FA) with 2% acetonitrile (ACN) prior to sample LC-MS analysis. Peptides were solubilized in 23µL of 0.1% aqueous FA with 2% ACN, adjusted to a concentration of 1µg/µL peptide, and an aliquot of retention time calibration peptides (i.e., 1 pmol/µL Biognosys iRT kit) was added in equal amounts to each sample prior to MS injection to correct for relative retention time differences between MS runs.

Data dependent mass spectrometry (DDA-MS) acquisition mode was used to inject the samples for the spectral library. DDA - LC-ESI-MS/MS was performed on a Q-Exactive HF Hybrid Quadrupole-Orbitrap Mass Spectrometer (Thermo Fisher Scientific) equipped with an Easy-nLC 1000 liquid chromatography system (Thermo Fisher Scientific). Peptides were trapped on an Acclaim pepmap100, C18, 3µm, 75µm x 20mm nano trap-column (Thermo Fisher Scientific) and separated on a 75 µm x 250 mm, C18, 2µm, 100 Å Easy-Spray column (Thermo Fisher Scientific). The analytical separation was run for 210 min using a gradient of H2O/FA 99.9%/0.1% (solvent A) and CH3CN/FA 99.9%/0.1% (solvent B). The gradient was applied at 250 nL/min by ramping solvent B from 4% to 23% in 160 min., then to 35% in 20 min., then to 90% in 10 min. This latter composition was maintained for 20 minutes. MS1 full scan resolution was set to 60’000 at *m/z* 200 with an AGC target of 3 x 106 and a maximum injection time of 60 ms. Mass range was set to 400-1250 *m/z.* For data dependent analysis (DDA), up to twenty precursor ions were isolated and fragmented by higher-energy collisional dissociation HCD at 27% NCE.

Resolution for MS2 scans was set to 15'000 at *m/z* 200 with an AGC target of 1 x 105 and a maximum injection time of 60 ms. Isolation width was set at 1.6 *m/z.* Full MS scans were acquired in profile mode whereas MS2 scans were acquired in centroid mode. Dynamic exclusion was of 20s.

Analysis of sample digests recorded using the Data Independent Mass Spectrometry (DIA-MS) acquisition mode for differential comparisons. DIA-MS DIA - LC-ESI-MS/MS was performed on a Q-Exactive HF Hybrid Quadrupole-Orbitrap Mass Spectrometer (Thermo Fisher Scientific) equipped with an Easy-nLC 1000 liquid chromatography system (Thermo Fisher Scientific). Peptides were trapped on an Acclaim pepmap100, C18, 3µm, 75µm x 20mm nano trap-column (Thermo Fisher Scientific) and separated on a 75 µm x 250 mm, C18, 2µm, 100 A Easy-Spray column (Thermo Fisher Scientific). The analytical separation was run for 150 min using a gradient of H2O/FA 99.9%/0.1% (solvent A) and CH3CN/FA 99.9%/0.1% (solvent B). The gradient was applied at 250 nL/min by ramping solvent B from 6% to 23% in 105 min., then to 35% in 20 min., then to 90%B in 10 min. with a final stay at this composition for 15 minutes. Data-Independant Acquisition (DIA) was performed with MS1 full scan at a resolution of 60,000 (FWHM) followed by 30 DIA MS2 scan with sequential fix isolation windows of 28 m/z covering a scan range from 400 to 1240 m/z. MS1 was performed in with an AGC target of 3 x 106, a maximum injection time of 60 ms and a scan range from 400 to 1240 m/z. DIA MS2 was performed using higher-energy collisional dissociation (HCD) at 27% with an AGC target of 1 x 106 and a maximum injection time of 50 ms.

Raw data (i.e., 96 raw DIA-MS files) were processed by commercial proteomic software package Spectronaut (version: 14.8.201029.47784, Biognosys, https://biognosys.com/software/spectronaut/). For differential analysis, Spectronaut DIA proteomics experiment was created. Quantitative data matrices for each sample type were generated with input of corresponding generated spectral library (HumanBrain3D_Neurotox.kit) and respective 96 DIA-MS raw data files. The proteins and peptide matrices were generated by default settings (i.e., BGS factory settings). Cross-run data normalization was performed on the whole data sets based on global median normalization of precursor intensities.

Normalized protein and peptide data were exported from Spectronaut software as csv. files for further analysis. Proteins not identified in the specific sample are annotated with NA (missing value). We have quantified nearly 6000 proteins in 90% of the samples. Technical replicates from selected samples display high positive correlation (Spearman's rho ≥ 0.94) across proteins quantified from the samples injected randomly across a measurement queue in the mass spectrometer.

MS files acquired in parallel in DDA mode (i.e., 31 raw MS files) were used to create a sample specific spectral library. For creation of spectral libraries we used default parameters against the ex_sp_9606_decoy.fasta database (the reviewed canonical Swiss-Prot complete proteome database for human, released 2014-01-24, entries 40’544) appended with common contaminants, reversed sequence decoys and iRT peptides. Included were trypsin digestion allowing two missed cleavages and ‘Carbamidomethyl (C)’ as static and ‘Oxidation (M)’ as variable modifications while minimum and maximum peptide length was set to seven and 52 amino acids, respectively. The mass tolerances were set to mode “dynamic” that is software determined tolerance based on extensive mass calibration and one-time correction factor was applied for precursor- and for fragment-ions. Unique proteins identified at 1% of protein false discovery rate (FDR) were included in libraries.

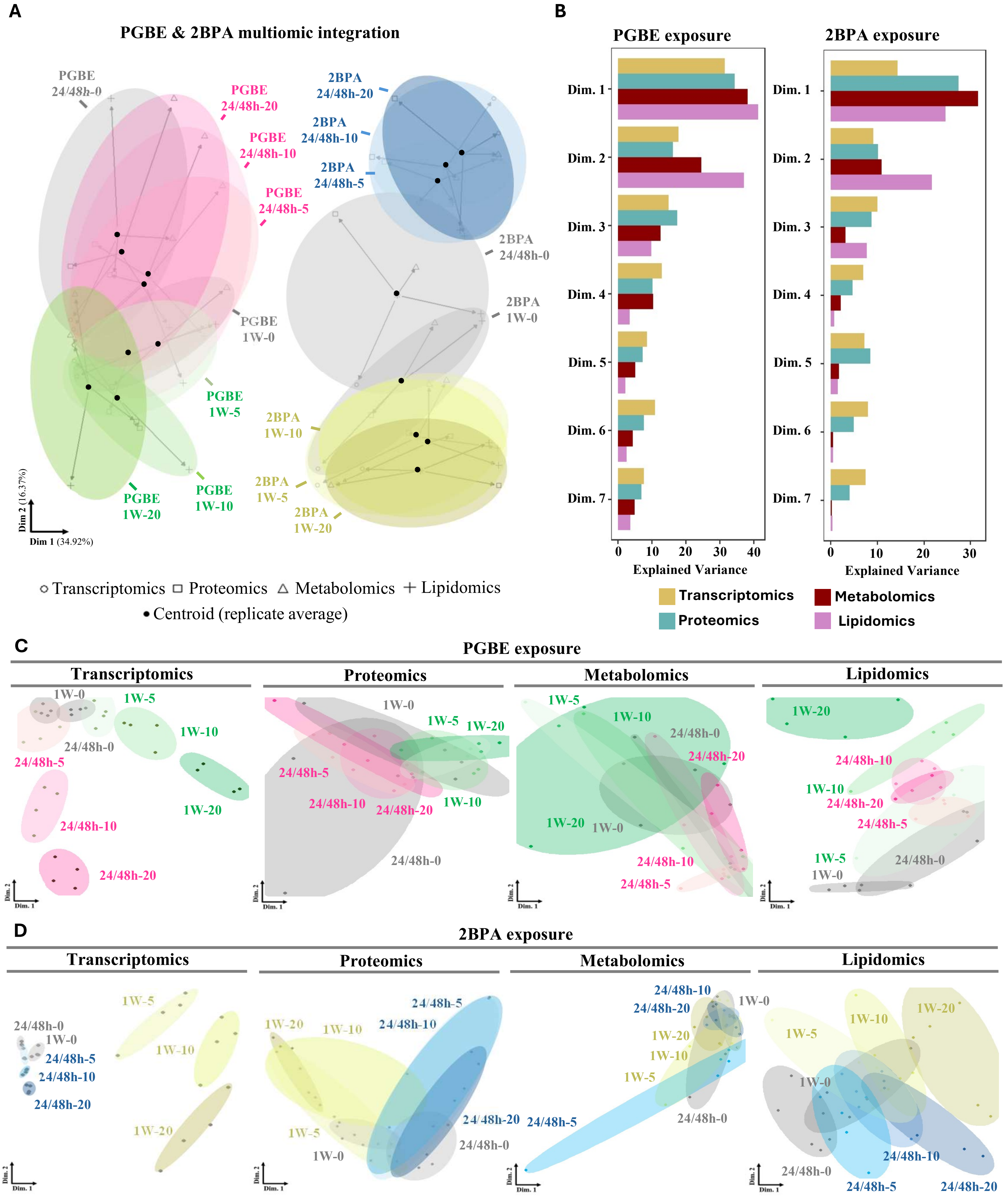

A
PGBE & 2BPA multiomic integration
PGBE 24/48h-0
PGBE 24/48h-20
PGBE 24/48h-10
PGBE 24/48h-5
PGBE 1W-0
PGBE 1W-5
PGBE 1W-10
PGBE 1W-20
2BPA 24/48h-20
2BPA 24/48h-10
2BPA 24/48h-5
2BPA 24/48h-0
2BPA 1W-0
2BPA 1W-10
2BPA 1W-5
2BPA 1W-20
Dim 2 (16.37%)
Dim 1 (34.92%)
○ Transcriptomics □ Proteomics △ Metabolomics + Lipidomics
• Centroid (replicate average)
B
PGBE exposure
2BPA exposure
Dim. 1
Dim. 2
Dim. 3
Dim. 4
Dim. 5
Dim. 6
Dim. 7
0 10 20 30 40
0 10 20 30
Explained Variance
Transcriptomics
Proteomics
Metabolomics
Lipidomics
C
PGBE exposure
Transcriptomics
Proteomics
Metabolomics
Lipidomics
1W-0
1W-5
1W-10
1W-20
24/48h-0
24/48h-5
24/48h-10
24/48h-20
Dim. 2
Dim. 1
D
2BPA exposure
Transcriptomics
Proteomics
Metabolomics
Lipidomics
1W-0
1W-5
1W-10
1W-20
24/48h-0
24/48h-5
24/48h-10
24/48h-20
Dim. 2
Dim. 1

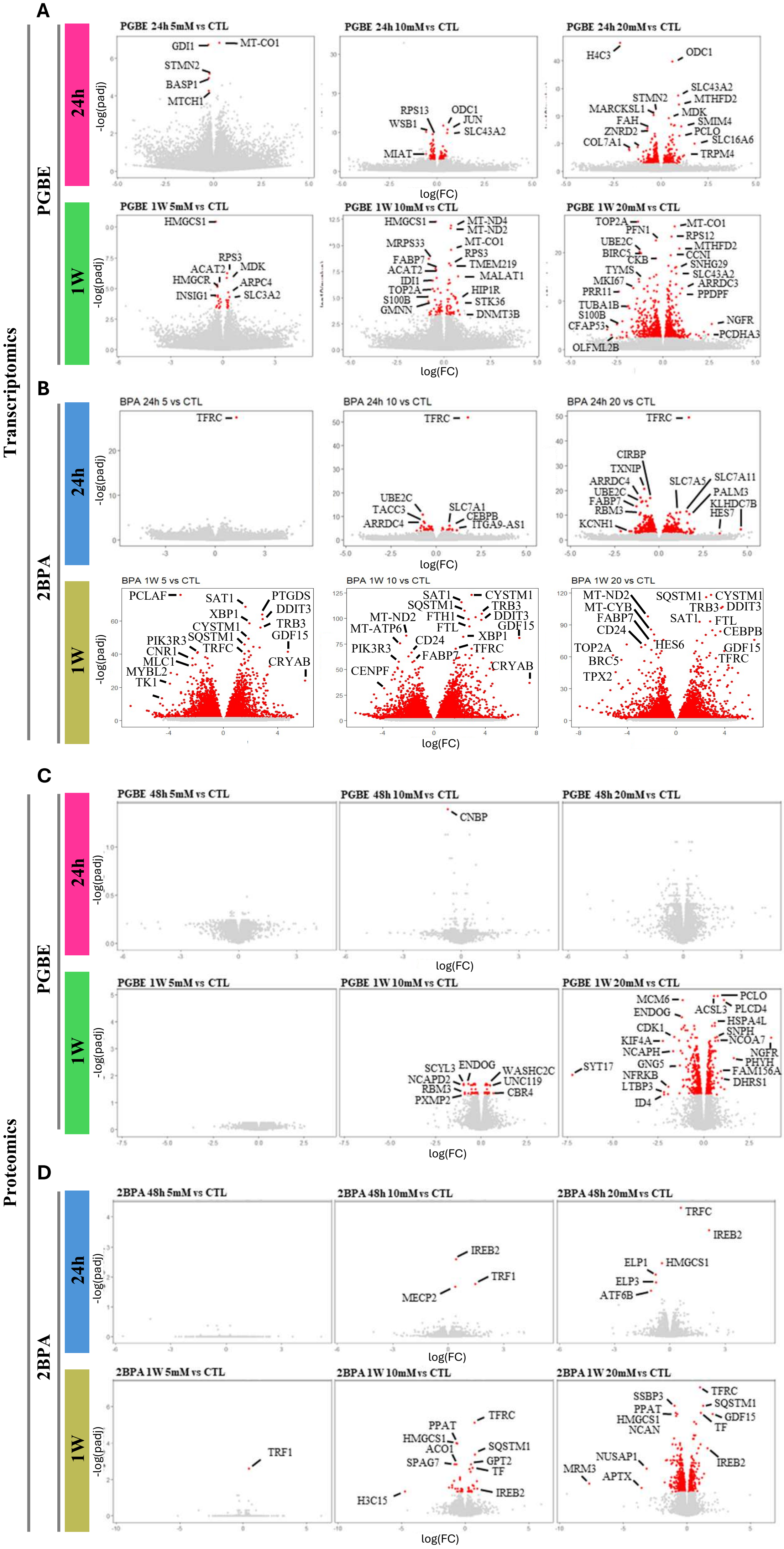
A
B
C
D
Transcriptomics
Proteomics
PGBE
2BPA
24h
1W
-log(padj)
log(FC)
PGBE 24h 5mM vs CTL
GDI1
MT-CO1
STMN2
BASP1
MTCH1
PGBE 24h 10mM vs CTL
RPS13
WSB1
ODC1
JUN
SLC43A2
MIAT
PGBE 24h 20mM vs CTL
H4C3
ODC1
STMN2
MARCKSL1
FAH
ZNRD2
COL7A1
SLC43A2
MTHFD2
MDK
SMIM4
PCLO
SLC16A6
TRPM4
PGBE 1W 5mM vs CTL
HMGCS1
RPS3
ACAT2
MDK
HMGCR
ARPC4
INSIG1
SLC3A2
PGBE 1W 10mM vs CTL
HMGCS1
MT-ND4
MT-ND2
MRPS33
MT-CO1
FABP7
RPS3
ACAT2
TMEM219
IDI1
MALAT1
TOP2A
HIP1R
S100B
STK36
GMNN
DNMT3B
PGBE 1W 20mM vs CTL
TOP2A
PFN1
MT-CO1
RPS12
UBE2C
MTHFD2
BIRC5
CKB
CCNI
SNHG29
TYMS
SLC43A2
MKI67
ARRDC3
PRR11
PPDPF
TUBA1B
S100B
NGFR
CFAP53
PCDHA3
OLFML2B
BPA 24h 5 vs CTL
TFRC
BPA 24h 10 vs CTL
TFRC
UBE2C
TACC3
ARRDC4
SLC7A1
CEBPB
ITGA9-AS1
BPA 24h 20 vs CTL
TFRC
CIRBP
TXNIP
ARRDC4
UBE2C
FABP7
RBM3
KCNH1
SLC7A5
SLC7A11
PALM3
KLHDC7B
HES7
BPA 1W 5 vs CTL
PCLAF
SAT1
PTGDS
XBP1
DDIT3
CYSTM1
TRB3
PIK3R3
SQSTM1
GDF15
CNR1
TRFC
MLC1
CRYAB
MYBL2
TK1
BPA 1W 10 vs CTL
SAT1
CYSTM1
SQSTM1
TRB3
FTH1
DDIT3
MT-ND2
FTL
GDF15
MT-ATP6
XBP1
CD24
PIK3R3
FABP7
TFRC
CENPF
CRYAB
BPA 1W 20 vs CTL
MT-ND2
SQSTM1
CYSTM1
MT-CYB
TRB3
DDIT3
FABP7
SAT1
FTL
CD24
CEBPB
HES6
TOP2A
GDF15
BRC5
TFRC
TPX2
PGBE 48h 5mM vs CTL
PGBE 48h 10mM vs CTL
CNBP
PGBE 48h 20mM vs CTL
PGBE 1W 5mM vs CTL
PGBE 1W 10mM vs CTL
SCYL3
ENDOG
WASHC2C
NCAPD2
UNC119
RBM3
CBR4
PXMP2
PGBE 1W 20mM vs CTL
MCM6
PCLO
ENDOG
ACSL3
PLCD4
CDK1
HSPA4L
KIF4A
SNPH
NCOA7
NCAPH
NGFR
SYT17
GNG5
PHYH
NFRKB
FAM156A
LTBP3
DHRS1
ID4
2BPA 48h 5mM vs CTL
2BPA 48h 10mM vs CTL
IREB2
TRF1
MECP2
2BPA 48h 20mM vs CTL
TRFC
IREB2
ELP1
HMGCS1
ELP3
ATF6B
2BPA 1W 5mM vs CTL
TRF1
2BPA 1W 10mM vs CTL
TFRC
PPAT
HMGCS1
ACO1
SQSTM1
SPAG7
GPT2
TF
IREB2
H3C15
2BPA 1W 20mM vs CTL
SSBP3
TFRC
PPAT
SQSTM1
HMGCS1
GDF15
NCAN
TF
NUSAP1
IREB2
MRM3
APTX

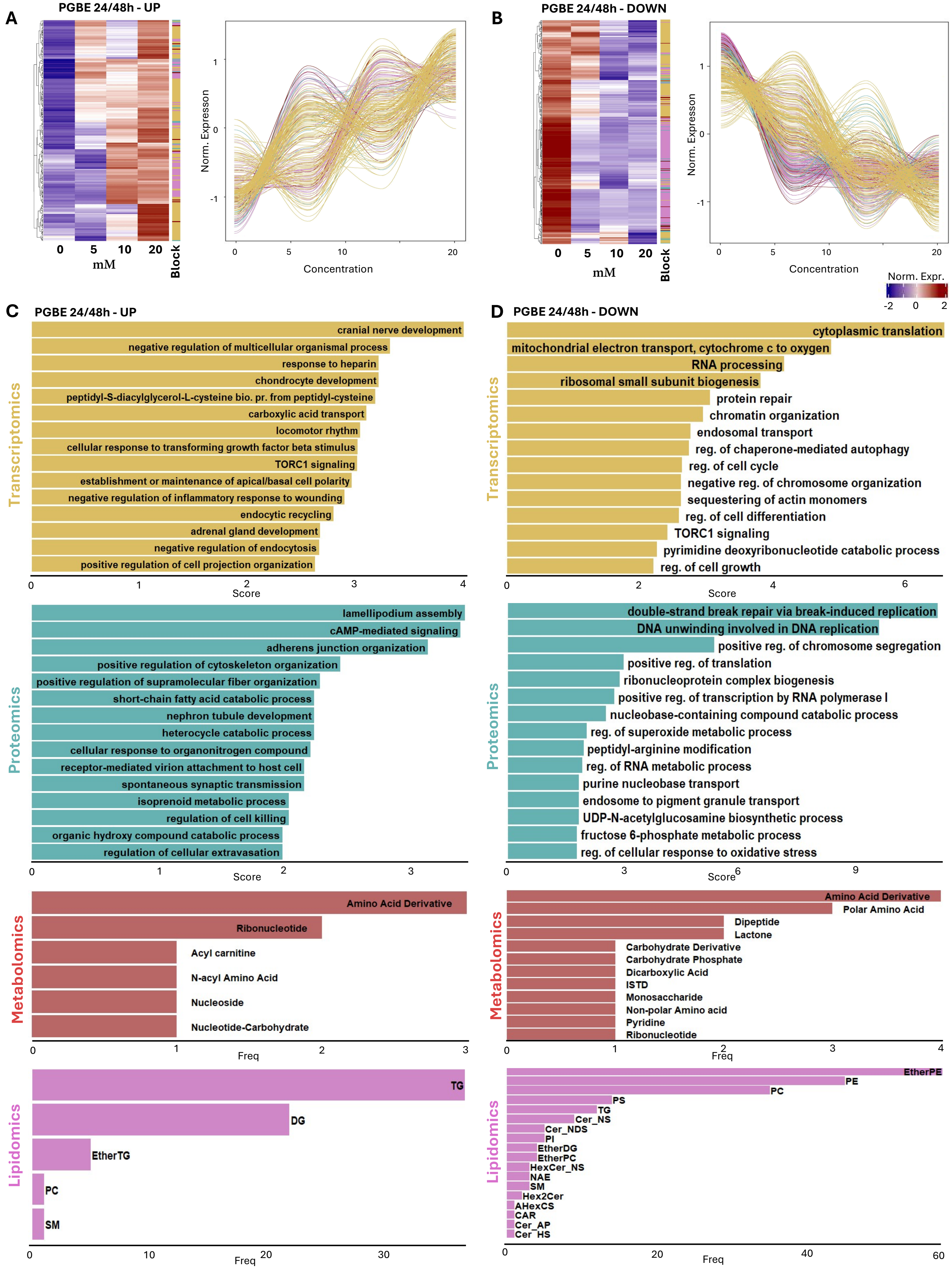
A
PGBE 24/48h - UP
0 5 10 20
mM
Block
Norm. Expresson
Concentration
B
PGBE 24/48h - DOWN
Norm. Expr.
-2 0 2
C
PGBE 24/48h - UP
Transcriptomics
cranial nerve development
negative regulation of multicellular organismal process
response to heparin
chondrocyte development
peptidyl-S-diacylglycerol-L-cysteine bio. pr. from peptidyl-cysteine
carboxylic acid transport
locomotor rhythm
cellular response to transforming growth factor beta stimulus
TORC1 signaling
establishment or maintenance of apical/basal cell polarity
negative regulation of inflammatory response to wounding
endocytic recycling
adrenal gland development
negative regulation of endocytosis
positive regulation of cell projection organization
Score
Proteomics
lamellipodium assembly
cAMP-mediated signaling
adherens junction organization
positive regulation of cytoskeleton organization
positive regulation of supramolecular fiber organization
short-chain fatty acid catabolic process
nephron tubule development
heterocycle catabolic process
cellular response to organonitrogen compound
receptor-mediated virion attachment to host cell
spontaneous synaptic transmission
isoprenoid metabolic process
regulation of cell killing
organic hydroxy compound catabolic process
regulation of cellular extravasation
Metabolomics
Amino Acid Derivative
Ribonucleotide
Acyl carnitine
N-acyl Amino Acid
Nucleoside
Nucleotide-Carbohydrate
Freq
Lipidomics
TG
DG
EtherTG
PC
SM
D
PGBE 24/48h - DOWN
cytoplasmic translation
mitochondrial electron transport, cytochrome c to oxygen
RNA processing
ribosomal small subunit biogenesis
protein repair
chromatin organization
endosomal transport
reg. of chaperone-mediated autophagy
reg. of cell cycle
negative reg. of chromosome organization
sequestering of actin monomers
reg. of cell differentiation
TORC1 signaling
pyrimidine deoxyribonucleotide catabolic process
reg. of cell growth
double-strand break repair via break-induced replication
DNA unwinding involved in DNA replication
positive reg. of chromosome segregation
positive reg. of translation
ribonucleoprotein complex biogenesis
positive reg. of transcription by RNA polymerase I
nucleobase-containing compound catabolic process
reg. of superoxide metabolic process
peptidyl-arginine modification
reg. of RNA metabolic process
purine nucleobase transport
endosome to pigment granule transport
UDP-N-acetylglucosamine biosynthetic process
fructose 6-phosphate metabolic process
reg. of cellular response to oxidative stress
Amino Acid Derivative
Polar Amino Acid
Dipeptide
Lactone
Carbohydrate Derivative
Carbohydrate Phosphate
Dicarboxylic Acid
ISTD
Monosaccharide
Non-polar Amino acid
Pyridine
Ribonucleotide
EtherPE
PE
PC
PS
TG
Cer_NS
Cer_NDS
PI
EtherDG
EtherPC
HexCer_NS
NAE
SM
Hex2Cer
AHexCS
CAR
Cer_AP
Cer_HS

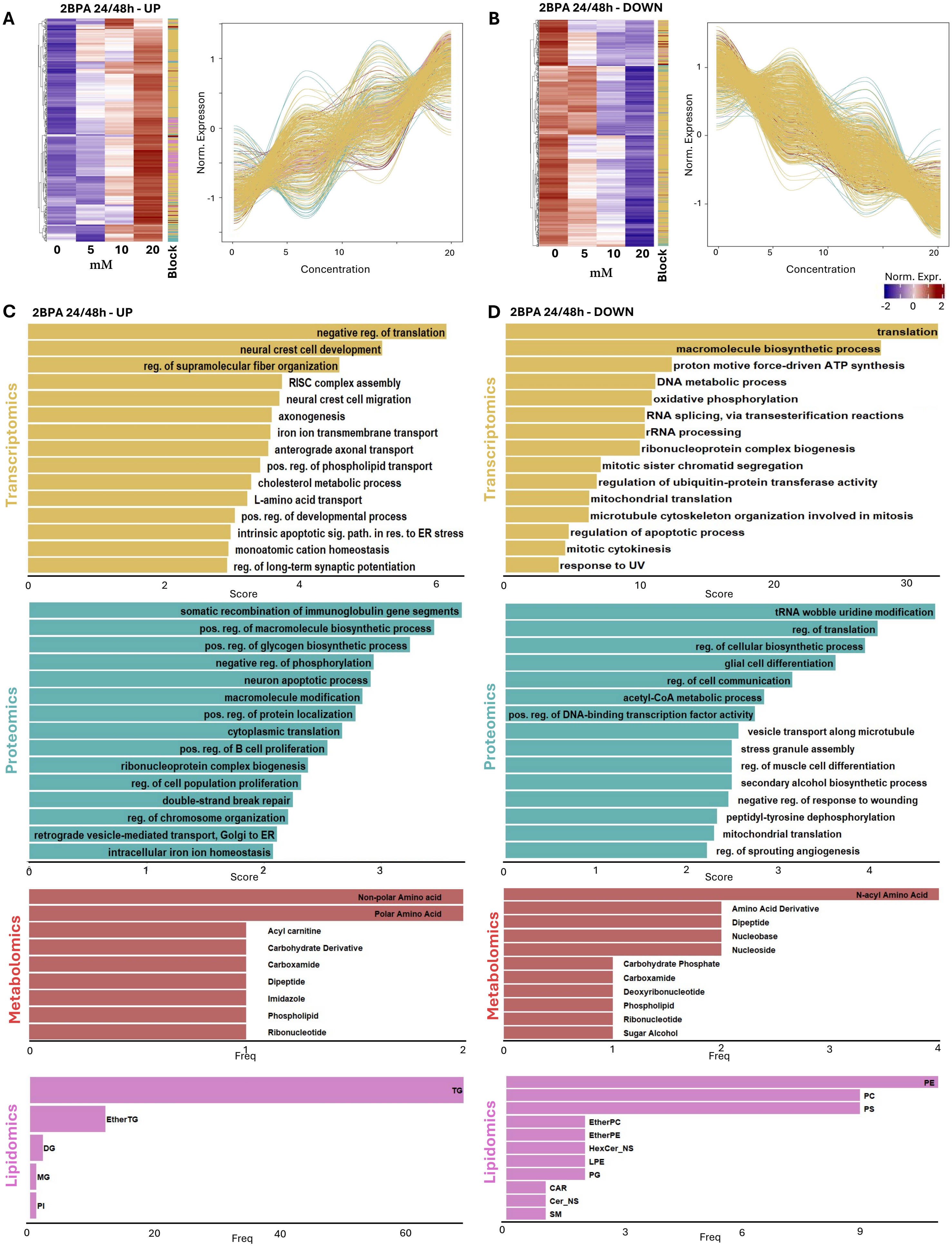

A
2BPA 24/48h - UP
0 5 10 20
mM
Block
Norm. Expresson
Concentration
B
2BPA 24/48h - DOWN
Norm. Expr.
C
2BPA 24/48h - UP
Transcriptomics
negative reg. of translation
neural crest cell development
reg. of supramolecular fiber organization
RISC complex assembly
neural crest cell migration
axonogenesis
iron ion transmembrane transport
anterograde axonal transport
pos. reg. of phospholipid transport
cholesterol metabolic process
L-amino acid transport
pos. reg. of developmental process
intrinsic apoptotic sig. path. in res. to ER stress
monoatomic cation homeostasis
reg. of long-term synaptic potentiation
Score
Proteomics
somatic recombination of immunoglobulin gene segments
pos. reg. of macromolecule biosynthetic process
pos. reg. of glycogen biosynthetic process
negative reg. of phosphorylation
neuron apoptotic process
macromolecule modification
pos. reg. of protein localization
cytoplasmic translation
pos. reg. of B cell proliferation
ribonucleoprotein complex biogenesis
reg. of cell population proliferation
double-strand break repair
reg. of chromosome organization
retrograde vesicle-mediated transport, Golgi to ER
intracellular iron ion homeostasis
Metabolomics
Non-polar Amino acid
Polar Amino Acid
Acyl carnitine
Carbohydrate Derivative
Carboxamide
Dipeptide
Imidazole
Phospholipid
Ribonucleotide
Freq
Lipidomics
TG
EtherTG
DG
MG
PI
D
2BPA 24/48h - DOWN
translation
macromolecule biosynthetic process
proton motive force-driven ATP synthesis
DNA metabolic process
oxidative phosphorylation
RNA splicing, via transesterification reactions
rRNA processing
ribonucleoprotein complex biogenesis
mitotic sister chromatid segregation
regulation of ubiquitin-protein transferase activity
mitochondrial translation
microtubule cytoskeleton organization involved in mitosis
regulation of apoptotic process
mitotic cytokinesis
response to UV
tRNA wobble uridine modification
reg. of translation
reg. of cellular biosynthetic process
glial cell differentiation
reg. of cell communication
acetyl-CoA metabolic process
pos. reg. of DNA-binding transcription factor activity
vesicle transport along microtubule
stress granule assembly
reg. of muscle cell differentiation
secondary alcohol biosynthetic process
negative reg. of response to wounding
peptidyl-tyrosine dephosphorylation
mitochondrial translation
reg. of sprouting angiogenesis
N-acyl Amino Acid
Amino Acid Derivative
Dipeptide
Nucleobase
Nucleoside
Carbohydrate Phosphate
Carboxamide
Deoxyribonucleotide
Phospholipid
Ribonucleotide
Sugar Alcohol
PE
PC
PS
EtherPC
EtherPE
HexCer_NS
LPE
PG
CAR
Cer_NS
SM

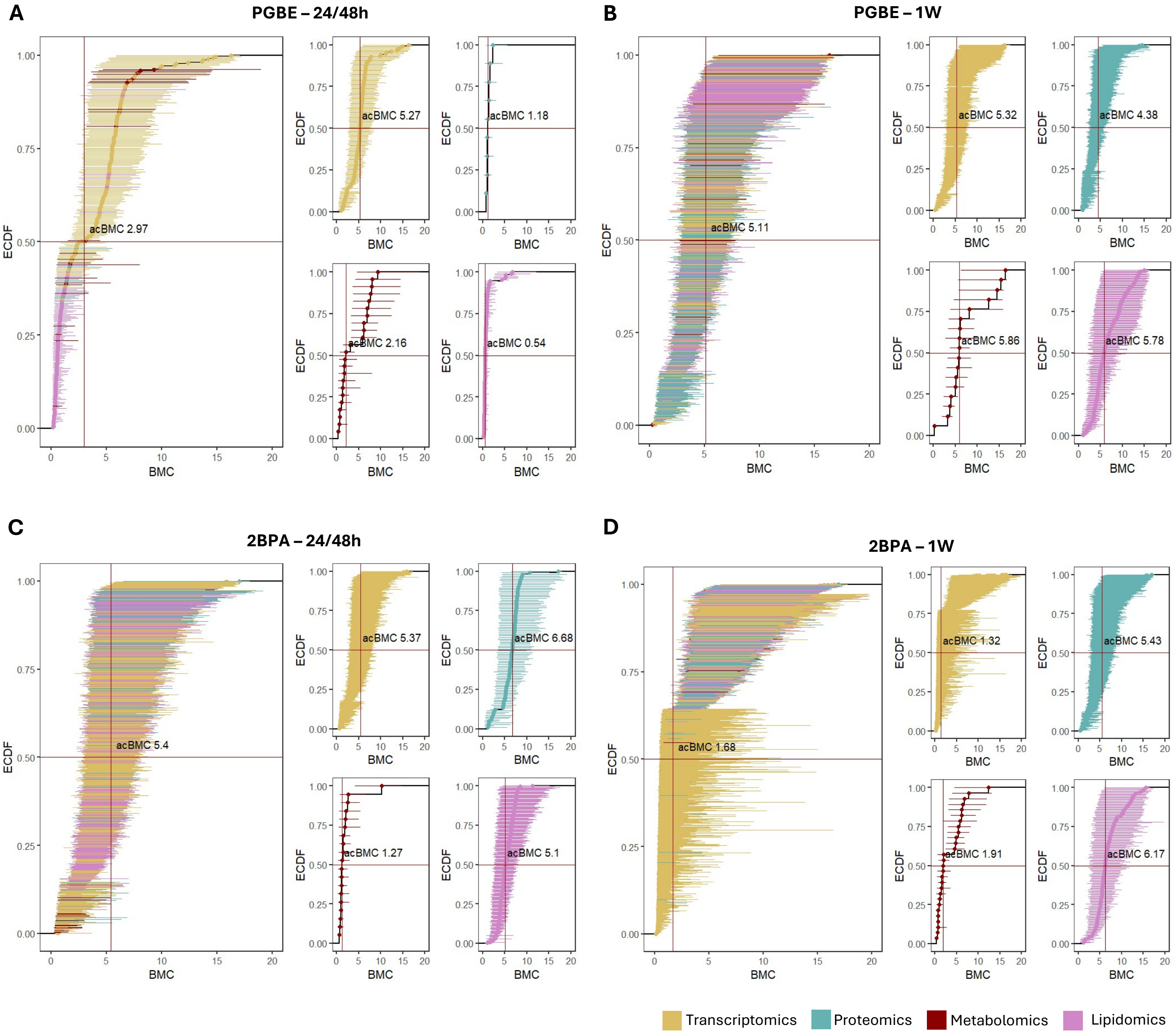
A
PGBE – 24/48h
B
PGBE – 1W
C
2BPA – 24/48h
D
2BPA – 1W
ECDF
BMC
acBMC 2.97
acBMC 5.27
acBMC 1.18
acBMC 2.16
acBMC 0.54
acBMC 5.11
acBMC 5.32
acBMC 4.38
acBMC 5.86
acBMC 5.78
acBMC 5.4
acBMC 5.37
acBMC 6.68
acBMC 1.27
acBMC 5.1
acBMC 1.68
acBMC 1.32
acBMC 5.43
acBMC 1.91
acBMC 6.17
Transcriptomics
Proteomics
Metabolomics
Lipidomics
